# Structural basis of lipid-mediated self-regulation of Tissue Factor

**DOI:** 10.64898/2026.09.07.749947

**Authors:** Alexei Iakhiaev

## Abstract

Tissue Factor (TF) is a tightly regulated transmembrane protein that maintains its encrypted state on the cell surface with an unknown conformation. Generation of TF conformational ensembles by the co-folding method using alphafold3 and boltz2 software revealed two mechanisms of TF self-regulation and corresponding conformational changes that were validated by Molecular Dynamics simulations. The first mechanism includes tilted and upright conformations of TF, whereas the second mechanism involves the spontaneous formation of TF oligomers, mostly dimers. The structural changes in TF responsible for both mechanisms were reversible and lipid-dependent. TF co-folding with phosphatidylserine resulted in approximately 90% of the TF extracellular domain in an upright conformation, enabling fast Factor VII (FVII) binding, while co-folding with phosphatidylcholine resulted in 60% of TF conformations tilted relative to the membrane surface and with a hidden FVII binding site. TF residues 210-219 (Linker Peptide) serve as a sensor that determines the TF conformation by interacting with the headgroups of phospholipids. TF self-association results in four types of homodimers with distinct relative orientations of extracellular domains and different accessibility of TF FVII binding sites. Approximately half of the dimers had accessible FVII binding sites, whereas in the remaining dimers, the FVII binding sites were partially or completely blocked. Co-folding of two TF sequences in the presence of cholesterol and phospholipids inhibited the dimerization and increased the number of TF monomers capable of FVII binding. These previously unknown TF conformations provide a structural basis for maintaining the encrypted and decrypted states of TF on the cell surface.

## INTRODUCTION

Tissue Factor (TF) is a type I transmembrane glycoprotein that initiates the extrinsic pathway of blood coagulation by forming a complex with the serine protease Factor VIIa (FVIIa). The TF-FVIIa complex activates Factor X (FX) to FXa, launching a downstream cascade of coagulation and anticoagulant reactions (1). Structural model of the full, membrane-bound ternary complex of TF-FVIIa-FXa is lacking at this time and the understanding of coagulation initiation machinery remains incomplete. This knowledge gap is related to the spatial arrangement of the components of the complex and the influence of the lipid bilayer on the conformation and orientation of the complex components (2, 3). Most importantly, many questions regarding the structural basis of the regulatory mechanisms of TF functional activity related to its encryption and decryption remain unanswered (4–7). The extracellular domain of TF (TFecd) binds FVIIa to activate the coagulation cascade, a transmembrane domain (TFtmd) anchors it in the cell membrane, and a short cytoplasmic domain (TFicd) composed of 20 amino acids involved in cell signaling (1). TFecd (residues 1–219) is composed of two fibronectin type III domains (N-terminal domain tagged as TFfn1 and membrane proximal domain TFfn2). TFecd binds to FVIIa, and the resulting TF-FVIIa complex binds to and activates coagulation factors X, IX, and VII, thereby initiating the blood coagulation cascade reactions.

The TFtmd domain (residues 220–242) anchors TF to the cell surface and plays a role in cell signaling pathways and regulation of TF activity. Membrane association is critical for TF function in coagulation. TFicd (residues 243–263) is non-essential for coagulation and plays a role in cell signaling. This domain contains sites for post-translational modifications, such as phosphorylation and cysteine palmitoylation, which regulate TF activity in response to stress (1).

Experimental structures of TFecd and TFicd have been published, whereas the TFtmd structure and full-length TF structures are available from modeling studies. Modeling studies, including MD simulations, have demonstrated that TFecd and TFtmd are connected by a charged and flexible linker peptide (LP), which allows the independent orientation of TFecd and TFtmd relative to the membrane bilayer and each other. The structure of LP is available from the AF3 predicted structural models database. This flexible peptide is located in the membrane-proximal region of TF and most likely interacts with the polar head of phospholipids. The conformation of this peptide can potentially regulate TF activity by changing the orientation of TFecd relative to the membrane surface, thereby generating a TF conformation that is more or less suitable for FVIIa binding to TF (3, 8).

Cell membrane phospholipids not only provide a platform but also serve as active determinants of whether TF remains a benign, encrypted molecule or becomes a powerful initiator of the blood clotting cascade. TF encryption refers to the suppression of procoagulant activity on cell surfaces, where it exists in a cryptic or inactive form. Decryption or activation occurs when TF can initiate blood coagulation. This process involves a complex interplay of factors, including changes in the cell membrane phospholipid composition and potential structural changes within the TF protein. Resting cells maintain an asymmetric distribution of phospholipids in the plasma membrane, with phosphatidylserine (PS) primarily located on the inner leaflet of the membrane. This asymmetry is disrupted upon cell activation, leading to PS exposure on the outer leaflet, which is crucial for TF activity (5, 9, 10). Increased intracellular calcium levels can trigger PS exposure and TF decryption. A PS-rich surface is crucial for the assembly and maximal efficiency of the TF-FVIIa complex. TF localization within lipid rafts may influence its encryption status (11–13). Lipid rafts are specialized regions of the plasma membrane enriched with cholesterol and sphingolipids, giving them a more ordered structure than the surrounding membrane. TF is often localized to or near lipid rafts. Studies suggest that lipid rafts are necessary for maintaining cellular TF in the inactive state (12–14). Disruption of these rafts can lead to the decryption and activation of TF procoagulant activities. Lipids can act as essential cofactors or molecular "glue," mediating the formation and stabilization of protein oligomers and multisubunit complexes. Specific lipids are required for the dimerization of certain transmembrane proteins. TF forms dimers or oligomers in the cell membrane; however, the selectivity of lipids for TF is unknown. Specific sphingolipids have also emerged as direct regulators of TF function. The enzyme Acid Sphingomyelinase (ASMase) hydrolyzes sphingomyelin (SM) into ceramide. The translocation of ASMase to the cell membrane and the resulting production of ceramide have been shown to trigger TF procoagulant activity in macrophages in response to inflammatory stimuli or viral infection. This mechanism appears to be independent of the PS exposure pathways (7, 15–17). Recent studies suggest that the oxidation state of a cysteine disulfide bond (Cys186-Cys209) within TF might be a key determinant of its activity, with the oxidized form (disulfide bond) being associated with active TF activity. Protein Disulfide Isomerase, an enzyme involved in disulfide bond formation and isomerization, may play a role in regulate TF activity by influencing the Cys186-Cys209 bond (13, 18, 19).

Controversies and unanswered questions. The exact molecular mechanisms underlying TF encryption and decryption, particularly the role of the Cys186-Cys209 disulfide bond, remain controversial. The formation of a C-terminal disulfide bridge (Cys186-Cys209) can lock TF in its active conformation, and the TF mutant p.C186S;p.C209S is known to decrease the affinity of FVIIa for TF. However, this mutant is not a perfect mimic of the cryptic state, as naturally cryptic TF retains its affinity for FVIIa. However, the precise interplay between phospholipid asymmetry, calcium signaling, and lipid raft association in the regulation of TF activity is not fully understood. This study provides novel insights into TF regulation; describes structural basis of two mechanisms of TF self-regulation and investigates molecular interactions behind these mechanisms. The first mechanism includes previously unrecognized tilted and upright conformations of TF determined by TF capability to sense phospholipid environment and adjust its conformation correspondingly. The second mechanism is the lipid-dependent self-association of TF, responsible for the formation of TF dimers with the Factor VII binding site blocked in some dimers. Cholesterol inhibits the dimerization of TF, resulting in the formation of monomeric TF with an open FVII binding site. These previously unrecognized TF conformations provide a plausible explanation for the experimental observations related to TF encryption and decryption.

## RESULTS AND DISCUSSION

### I.1 TF Conformations

The structure of the extracellular domain of TF (amino acid residues 1-212) has been resolved experimentally. The structure of flTF is known from modeling studies only. The flTF structure (panel A) predicted by the AF3 software demonstrated that the structure of the region between TFecd and TFtmd was colored as a low-confidence, flexible, or disordered structure. This is consistent with previous observations that TFecd and TFtmd behave as independent structural elements that can change their relative orientation. The TF-FVIIa complex is oriented in a direction perpendicular to the membrane surface (Fig. 1B). Models of TF generated in the presence of water molecules or without additional molecules (Fig. 1 panels C and D) demonstrated a TF conformation where TFecd was tilted and could form different angles up to 90 °relative to the normal to the membrane surface plane. Because the optimal binding conditions for FVIIa require TFfn2 orientation in the direction close to the membrane normal, tilting TF leads to conformations with different angles relative to the membrane surface, creating the possibility of a previously unrecognized mechanism of TF regulation. As seen from Fig.1C, most of tilted conformations were hindering FVII binding to TF by hiding the FVII binding face of TF. However, tilt angles were different and it was possible to find tilted TFecd domain with open FVII binding face of TF (Fig. 1D). One could expect that the rate of FVIIa binding to TF tilted toward the membrane surface could be slower than the rate of binding to TF oriented optimally for FVIIa binding. This may result in inactive or diminished TF activity in the initiation of coagulation, which is a hallmark of encrypted TF. The possibility of the existence of a TF conformation corresponding to the encrypted TF warranted a detailed computational study to reveal the conditions that support this conformation and the molecular mechanism leading to the acquisition of this conformation by TF.

**Figure 1.**
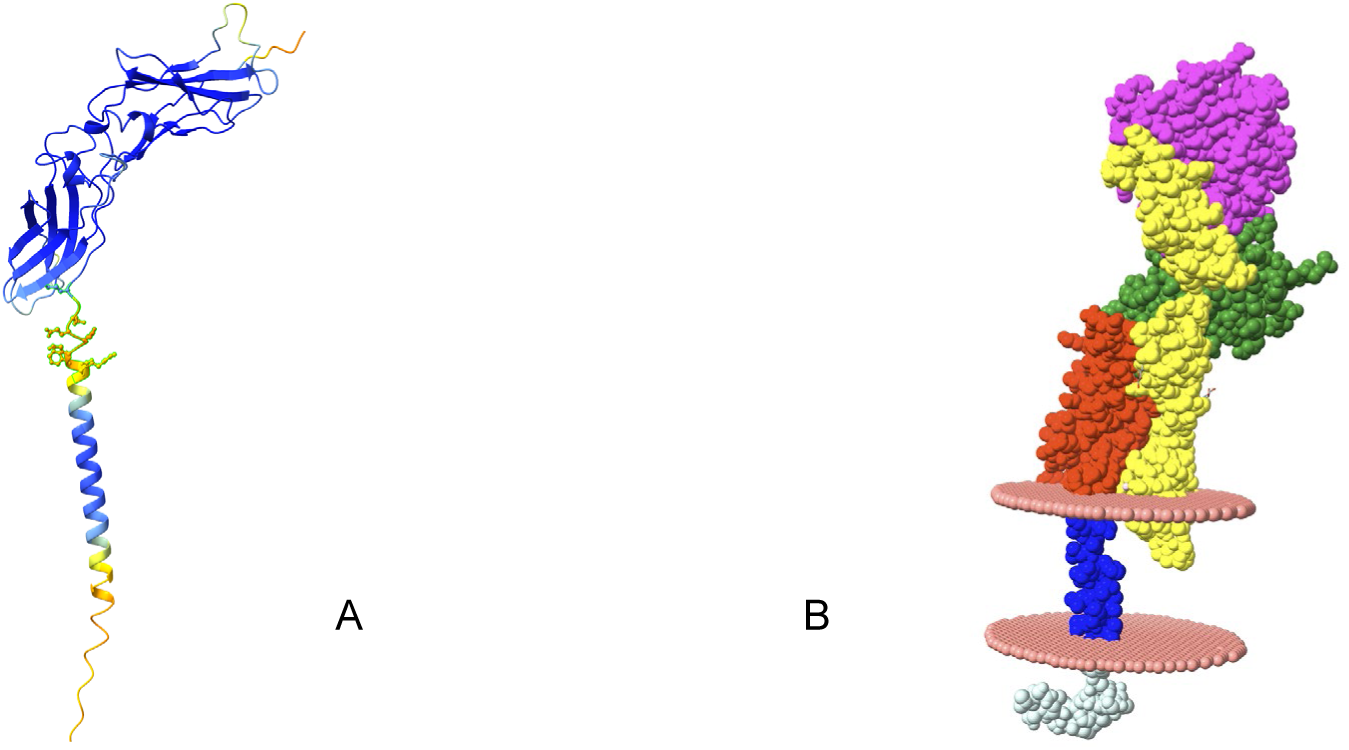
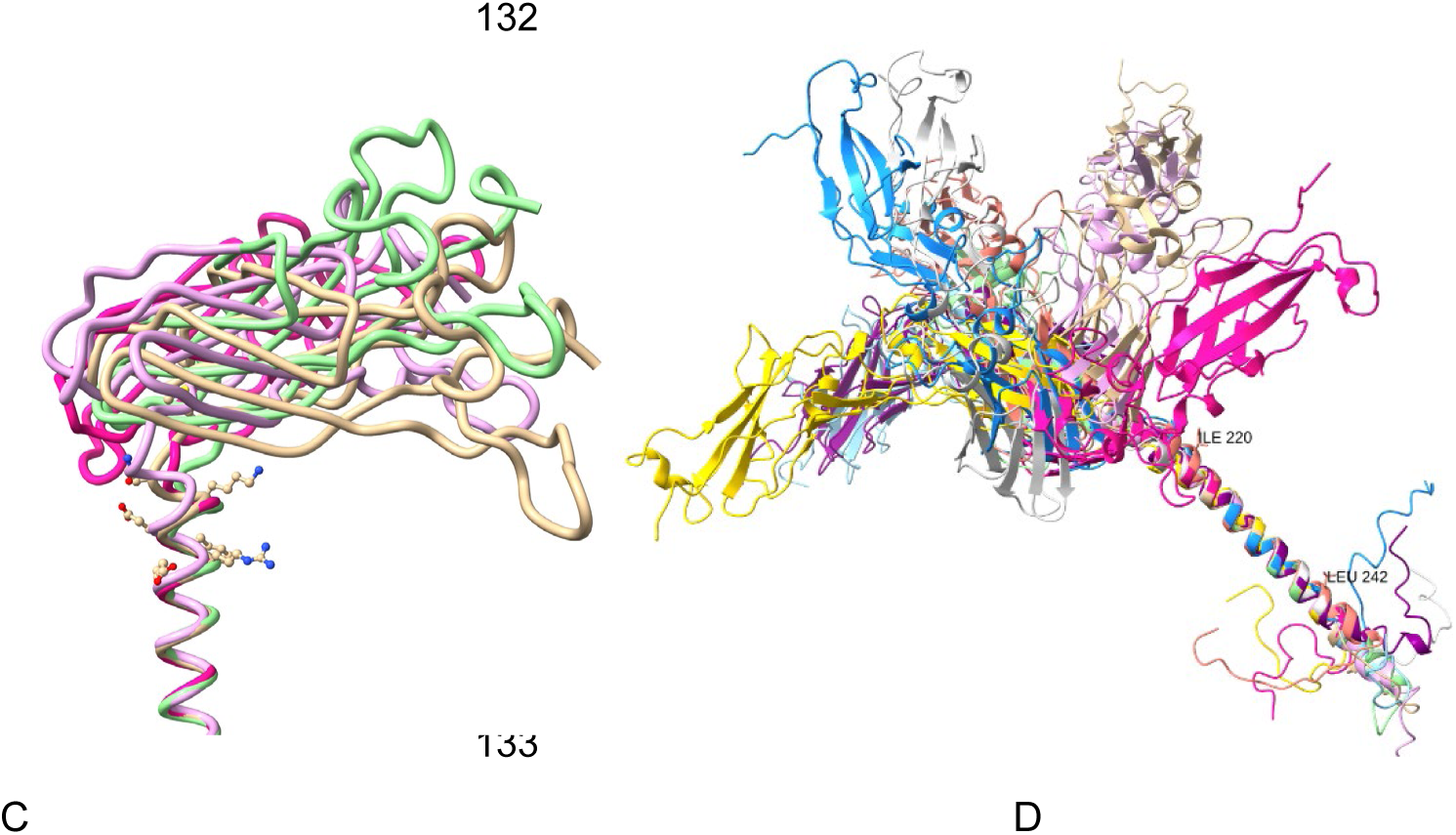
Structures of full-length TF and TF-FVIIa complex inserted into the lipid bilayer. Panel A. AF3 model of full-length TF (canonical amino acid sequence) colored by model confidence score and presented as a cartoon. Meaning of colors: Dark Blue, very high confidence; Light Blue/Green, from confident to medium-high confidence; yellow, low confidence, flexible or disordered structure; red, very low confidence. The ten amino acid-long LP connecting TFecd and TFtmd is shown as sticks and balls. Panel B. Space-filled representation of TF-FVIIa complex in the membrane colored by domains or chains. The image was generated using the structure from the file with the PDB ID 1DAN.pdb by superimposing the full-length TF and adding lipids. TF fibronectin type III domain 1 (TFfn1) is green, TFfn2 is red, and TFtmd is blue. FVIIa light chain, yellow; FVIIa heavy chain, magenta. The membrane bilayer is presented, showing the positions of the phospholipid polar headgroups as dummy-atoms. Panel C. Predicted structures of TF membrane-proximal domains, including TFfn2-TFtmd, in vacuum. The models were superimposed by matching the TFtmd domains, and each model was highlighted with a different color. Ten amino acids long linker peptide is shown as sticks and balls for model 1. Panel D. Predicted conformations of flTF, in vacuum without water or lipid molecules. The models were superimposed by matching the TFtmd domains, and each model was highlighted with a different color. The first (Ile220) and last (Leu242) amino acid residues of TFtmd are labeled.

**Figure 2.**
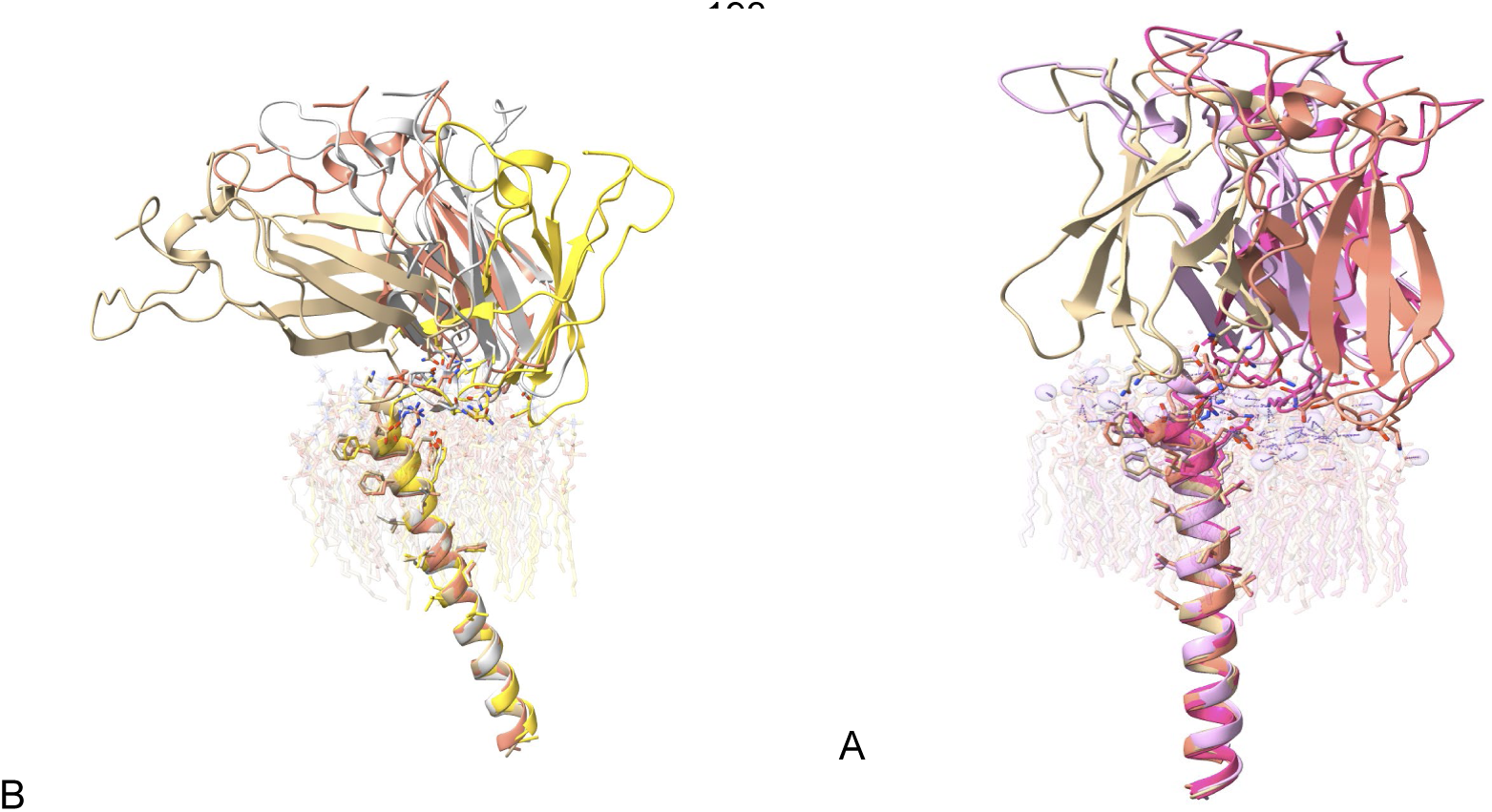
Structures of TFfn2_TFtmd co-folded in the presence of phospholipids DLPC and DLPS. Panel A. Boltz model of TFfn2-TFtmd co-folded in presence of phospholipids DLPC. Four model outputs colored separately were superimposed using their TFtmd domains. Phospholipid DLPC (16 molecules per model) is shown with 90 % opacity. Panel B. Boltz model of TFfn2-TFtmd co-folded in presence of 16 phospholipid DLPS molecules. Four individually colored model outputs were superimposed using their TFtmd domains. Phospholipid DLPS (16 molecules per protein) is shown with 90 % transparency.

### I.2 Co-folding experiments demonstrated dependence of TF conformation from the phospholipid environment

Co-folding, which utilizes generative models implemented in Boltz2 and AF3, represents the next generation of molecular docking suitable for modeling flexible binding sites. Co-folding can handle conformational changes in proteins during ligand binding, making it the method of choice for studying the phospholipid dependence of TF conformations. Study of conformational ensemble of both flTF and its membrane-proximal fragment (TFfn2-TFtmd-TFicd domains) demonstrated that TF is best described by a model composed of domain-flexible linker-domain. Two domains (TFecd and TFtmd) can be represented by rigid bodies while TF210-219 sequence as a flexible LP. The prediction of the distribution of the poses of rigid domains connected by flexible linkers remains an unsolved problem for machine learning predictions of the final structure (20). This system can be described using a population-weighted ensemble of discrete structures rather than a single average structure. The simplified representation we used describes conformation ensembles by interdomain angles and expected fast or slow rates of FVIIa binding to TF depending on the domain tilt relative to the membrane surface.

#### I.2.1. TF folded in the absence of phospholipids has a tilted conformation

The principal axis of TFfn2 was tilted by approximately 90 °relative to the principal axis of TFtmd (Fig. 1C). This was observed in the folding of the TF sequence alone or in the presence of molecules that did not interact with the membrane-proximal amino acids of TF. The flTF downloaded from the AF3 database had an angle between the principal axes of TFfn2 and TFtmd of approximately 10 degrees (Fig. 1A). For the TF structure folded in the presence of water or without any ligand, the TFfn2-TFtmd angle was 90+-20 degrees (Fig.1C and 1D). TF folded in the presence of 10 molecules of myristic acid (14:0) or palmitic acid (16:0) tilted to approximately the same extent as TF folded without ligands. Similar results were obtained for short-chain PSF when the number of molecules was less than 10, and lipid bilayer was not formed.

#### I.2.2. Phospholipid bilayer effects TF folding and the effect depended on the type of phospholipid

In the presence of DLPE, the TFfn2-TFtmd angle was also 90 ± 20 degrees (Fig.1C), and the interaction of DLPE with the LP was not detected. Co-folding with DLPC resulted In the TFfn2-TFtmd angle of 20 ± 20 degrees (Fig.1C). When the number of DLPC molecules (10–20) was sufficient to form a structure resembling a monolayer and the LP was buried in the polar headgroups of phospholipids, the number of molecules that could form a TF-FVIIa complex without clashes was approximately 70 % (seven out of ten structures). Some of these structures were tilted, suggesting a slower rate of FVIIa binding to the encrypted TF. Structures with clashes are considered inactive or potentially slower FVIIa binders than tilted TF. The average of several co-folding predictions was about 50 % of tilted structures in DLPC monolayer with TFfn2-TFtmd angles that were not favorable for FVIIa binding. The conclusion about FVIIa binding was based on sterical hindrance, which tilted conformation creates for FVIIa during accessing the TF residues involved in FVIIa binding. Approximately 90 % of the TF models generated in the presence of DLPS had an upright conformation, which should be capable of binding FVIIa at rates higher than those of the tilted conformations. Visual and Pylipid-assisted analyses of the contacts between phospholipids and TF amino acid residues demonstrated the prevailing interactions of DLPS and DLPC polar head interactions with LP amino acids KE-FRE with DLPC, Q--EFRE, and K214 with DLPS. Other interactions were observed with the TFtmd domain and two amino acids from the membrane-proximal loop of TF.

Co-folding experiments in the presence of PSF, DLPS, and PCPS phosphatidylserine species resulted in TF molecules that did not clash with FVIIa when superimposed with TF-FVIIa structure. Fewer tilted molecules were observed among these structures than among TF co-folded in the presence of PC or PE. Similarly, co-folding of the TF sequence in the presence of phosphatidic acid resulted in TF conformations that did not create clashes and with upright conformation of TFecd. In summary, TF folding in the presence of negatively charged PS and phosphatidic acid resulted in TF conformations that corresponded to active conformations favorable for FVIIa binding in approximately 90-100 % of the models.

### I.3. The MD study validated the phospholipid-dependence of TF conformations

The results of the co-folding studies demonstrated that both PC and PS polar headgroups interact with TF LP and induce different conformations characterized by different angles between the principal axes of TFfn2 and TFtmd domains. Although less accurate than the axis-vector representation of the relative orientation of these domains, the calculated angle allowed us to describe the TFecd conformation ensemble. A dataset that used backbone dihedral angles as features and interdomain angles as prediction targets enabled the use of RF regression to predict this angle and calculate the importance of different LP residues for phospholipid dependent conformations. The main goal for MD study was to visualize and quantify the structural changes in TF induced by lipids, which cannot be easily captured by traditional experimental methods. Because co-folding experiments demonstrated that TF LP could represent the proposed phospholipid sensor that induces conformational changes, we focused on TFfn2 (C-terminal membrane proximal fibronectin type III domain of TF) connected to TFtmd via LP. The presented results were obtained using the juxtamembrane domains embedded in DLPC, DLPS, and their mixtures. Similar results were generated for flTF as well (not shown).

#### I.3.1. MD simulation of TFfn2-TFtmd in membrane with different starting positions of LP relative to polar headgroups of DLPS

TFtmd α-helix embedded in DLPS bilayer experiences a hydrophobic mismatch because thickness of bilayer is smaller than the TFtmd length along its principal axis. Therefore, embedded TFtmd is oriented in an angle relative Z-axis (direction of the normal to the bilayer surface) and TFfn2 is tilted toward the membrane with LP amino acid residues contacting DLPS polar headgroups (Fig. 3A). MD simulations with these starting orientations in the presence of negatively charged PS headgroups induced a significant change in the tilt angle, leading to upright position TFfn2 relative to the membrane surface. At approximately 100 ns of simulation, TFfn2 was almost perpendicular to the membrane surface, which is the optimal TF binding conformation for FVIIa (Fig. 3B). This change was induced by interaction of DLPS polar headgroups with LP. During 400 ns long MD simulation, we observed two more conformational changes, when upright TFecd position become tilted again and finally, it returned back to the upright conformation. The conformational changes coincided with lost or establishment of contacts between LP and DLPS. These data suggest that the conformational changes of TF related to the LP region are fast (require less than 100 ns) and reversible (two cycles of conformational changes were observed during the 400 ns simulation). These conformational changes at the membrane interface alter the average height and orientation of TFecd relative to the membrane surface. These conformations were related to the contacts of LP amino acid residues with DLPS polar headgroups: contacts lead to upright conformation, while loss of contacts to tilted conformation. Specific TF-PS interactions involved in these conformational changes were LP Lys-Arg residues that can form stable salt bridges or hydrogen bonds with the negatively charged headgroups of PS lipids.

**Figure 3.**
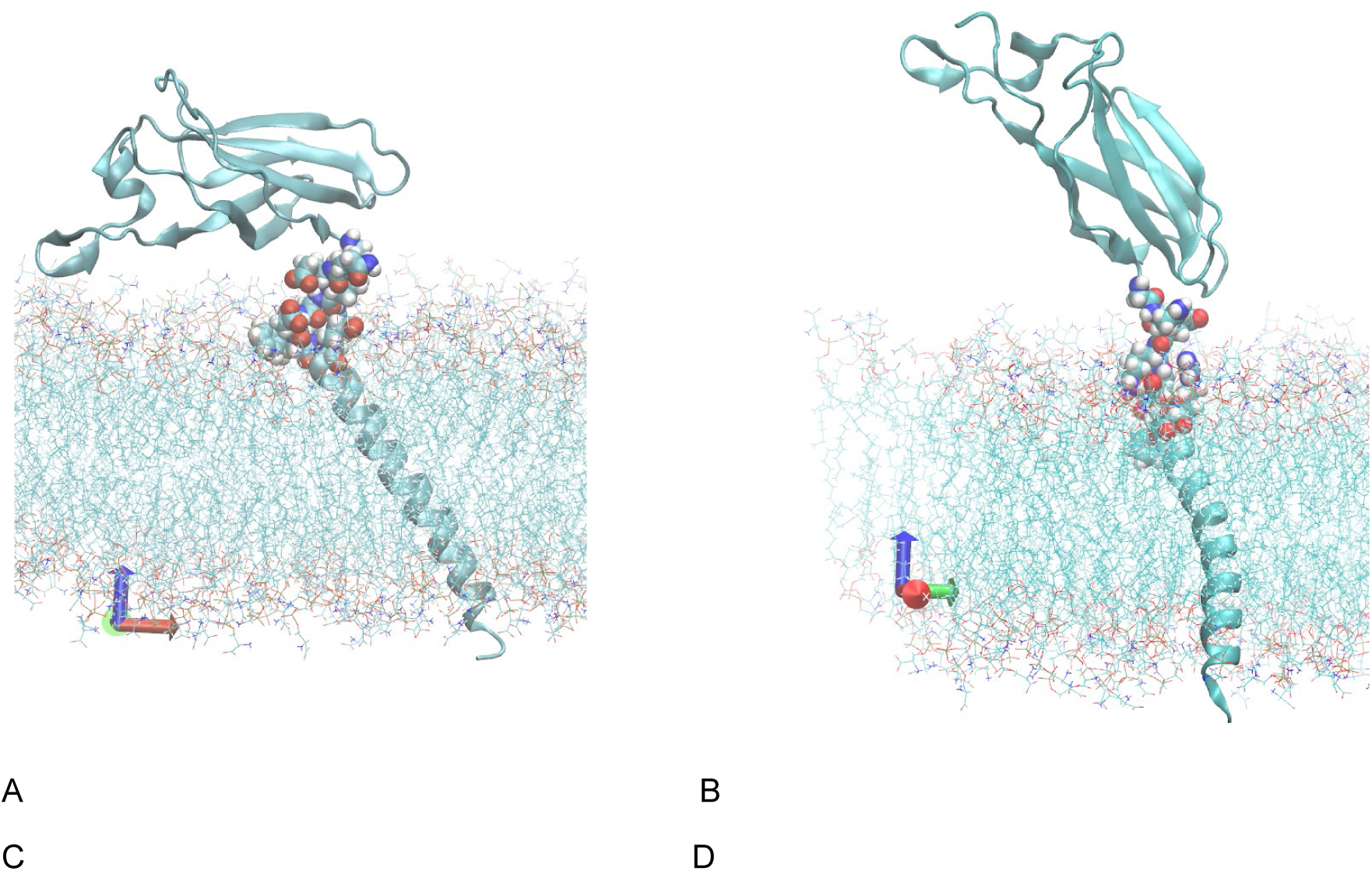
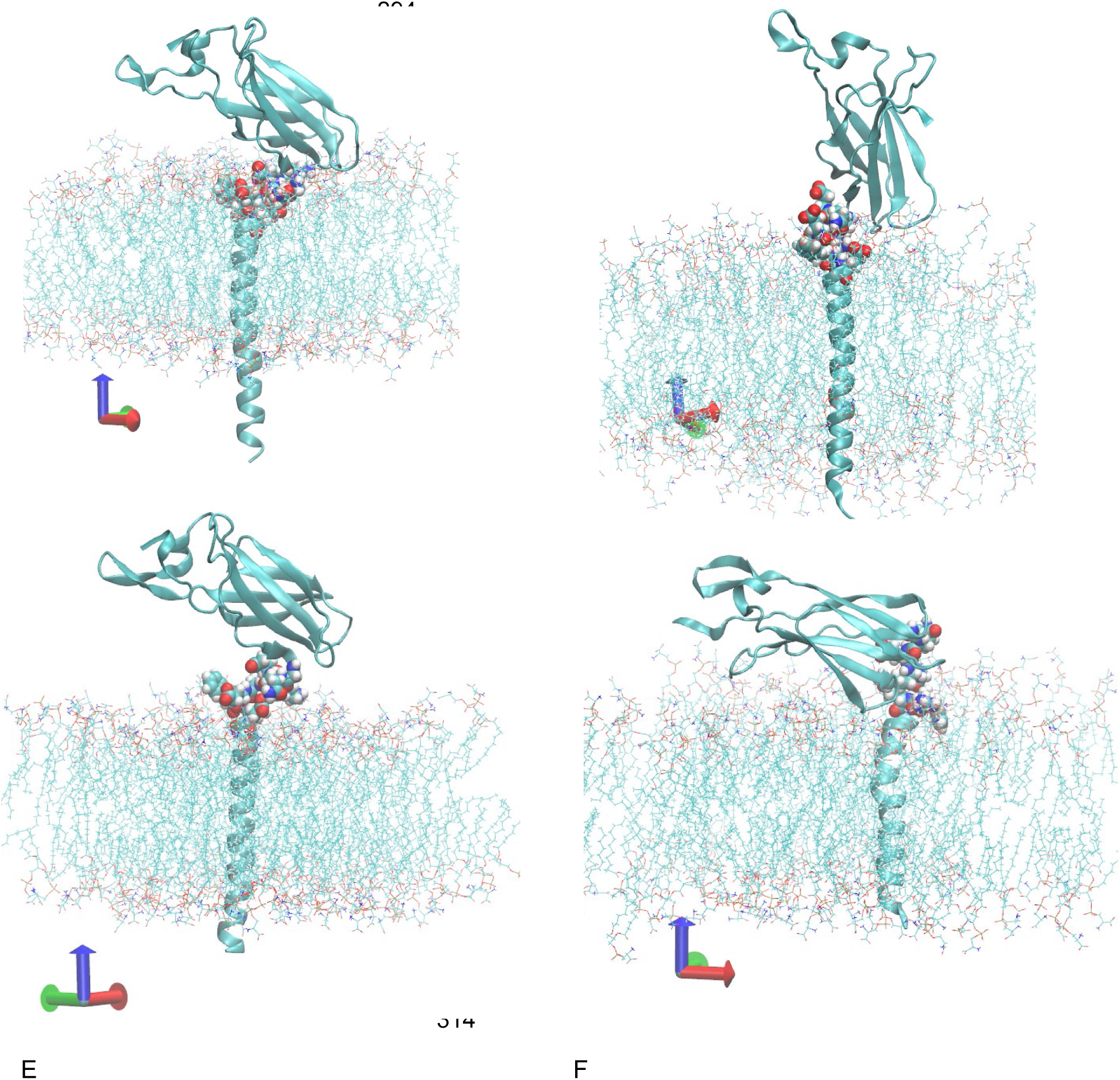
MD simulation of membrane proximal TFfn2-TFtmd domains embedded in the DLPS membrane bilayer in three different starting positions of LP relative to the membrane surface. Panel A. Tilted orientation of the fragment suggested by the OMP server at the beginning of simulation (at 0 ns run). Amino acid residues of LP peptide are shown as VMD VDW representation, protein is shown as cartoon, and DLPC molecules are in line representation in all panels. Panel B. Tilted orientation of TFfn2 changes to an orientation with approximately collinear position of the principal axis of TFtmd and TFfn2. The angle between the normal to the membrane surface was much smaller than the initial angle for TFfn2 and could reach approximately perpendicular to the membrane surface orientation after 150 ns of the production run of the MD simulation. Panel C. Tilted TFfn2 fragment with buried LP at the beginning of the simulation ( 0 ns run) with TFtmd approximately perpendicular to the membrane surface. Panel D. Fast (within 80 ns) reorientation of TFfn2 for buried LP interacting with polar headgroups of DLPS. Panel E. Perpendicular to the membrane surface orientation of TFtmd and tilted TFfn2 in the MD simulation, similar to Panels C and D, except without LP contacts with DLPC polar headgroups (TFfn2 membrane complex structure at 0 ns MD simulation). Panel F. Tilted orientation of TFfn2 in the simulation without LP contacts with the DLPS polar headgroups remained tilted even after the 500 ns MD simulation.

#### I.3.2. TFfn2-TFtmd in DLPS when TFtmd was perpendicular to the membrane surface

To verify the role of LP contacts with DLPS in acquiring the upright conformation by the TFfn2, the MD simulation of the system described in the previous section was repeated with changes of the initial orientation of TFtmd (Fig 3, panels C and D). In this simulation, the orientation of TFtmd in DLPS bilayer was along Z-axis, while TFfn2 retained approximately 90° angle to the TFtmd principal axis. The C245 residue in these experiments was modified to CYSP (palmitoylated cysteine) to help TFtmd maintain its orientation despite the hydrophobic mismatch. This resulted in the loss of contacts between the DLPS polar headgroups and LP and increased distance of TFfn2 from bilayer. TFecd reorientation to upright conformation was not observed during the 550 ns simulation that started from these initial conditions, whereas small change in the TFtmd angle relative to the membrane plane was observed. This observation supported the requirement of direct interaction between the phospholipid polar headgroups and LP residues to induce conformational changes, leading to domain reorientation.

#### I.3.3. TFfn2-TFtmd in DLPS when TFtmd is perpendicular to the membrane surface and the molecule is buried by 10 Å

To further elucidate the mechanism of phospholipid environment sensing by the TF LP, we simulated the same system described in the above section after shifting the entire TFfn2- TFtmd complex by 10 Å along the Z-axis to enable contact of the TF LP with the DLPS polar headgroups (Fig 3 panels E and F). Consistent with the observations from the co-folding experiments and MD simulations, the initially tilted TFecd changed its orientation to approximately perpendicular to the membrane surface orientation within approximately 100 ns of the simulation. This simulation continued after TFtmd moved along the Z-axis and lost LP residues contacts with the DLPS polar headgroups, the orientation of TFecd changed back to tilted and after some fluctuations returned back to orientation perpendicular to the membrane surface. These observations confirmed the requirement of LP-DLPS interactions for TF conformational changes.

#### I.3.4. MD simulation of TFfn2_TFtmd in the DLPC bilayer

The TF conformation, which was initially perpendicular to the membrane surface, acquired a tilted conformation at the end of the simulation of TFfn2 embedded in DLPC (data not shown). This observation was consistent with the co-folding experiments performed in the presence of DLPC.

The Cys245 residue was modified by palmitoylation (renamed as CysP) to facilitate TFtmd movement along the Z-axis to balance the hydrophobic mismatch between the TFtmd length and the thin bilayer formed by DLPC or DLPS which contain 16 carbon fatty acids. In a tilted TFtmd, the mismatch was balanced by tilting the TFtmd. When TFtmd orientation was perpendicular to the membrane plane, the length of TFtmd was longer than the DLPS membrane thickness, and the direct contacts between LP and polar headgroups of DLPS were removed. The 10 Å shift along the Z-axis allowed the reestablishment of LP contacts. The effect of CysP, together with the balancing of the hydrophobic mismatch, allowed us to study the consequences of the loss of direct contacts between LP and the polar headgroups of phospholipids. The loss of contacts owing to the shift of TFtmd along the Z-axis was observed within 100 ns.

### I.4. LP as a lipid sensor. Mechanistic explanation of lipid sensing by TF

Co-folding experiments and MD simulations suggest that the contacts of LP amino acid residues with polar headgroups of phospholipids determine tilted or upright orientation of TFecd relative to the membrane surface, which correlates with preferred or unfavorable conformations for binding of FVIIa to TF. To elucidate the mechanism of phospholipid sensing by TF LP, we conducted an extensive study of the TF conformations resulting from the co-folding of TF mutants in the presence of DLPS and DLPC liposomes. These studies included mutations in the LP to replace charged and polar amino acid residues with alanine or glycine, introduction of additional glycine hinges into LP, replacement of two glycine residues in wild-type LP with Valine and Isoleucine to stiffen LP, and breaking mutations G211P and G215P. As a baseline for comparisons 50 % and 90 % upright conformations for wild-type TF in DLPC and DLPS bilayers, respectively were used.

#### I.4.1. Co-folding of the TF mutant, where amino acids 212-QEKGEFRE-220 of LP were replaced with eight alanine residues

Co-folding of the TFfn2-TFtmd sequence in the presence of eight DLPC and eight DLPS phospholipid molecules resulted in 100 % tilted molecules with an angle of approximately 90 °between the TFfn2 and TFtmd domains. Approximately 80 % of these molecules demonstrated a possible FVIIa clash with the TF LP peptide and the amino acids at the beginning of the TFtmd. These results are consistent with the hypothesis that the charged amino acid residues of LP are involved in sensing the lipid environment and changing the TF conformation depending on the phospholipid environment. The results in the presence of 16 phospholipid molecules were similar to those described above; 85 % of the conformations in the presence of DLPS and 95 % in the presence of DLPC could be classified as inactive.

#### I.4.2. Co-folding of LP mutants, where two neighboring amino acids were replaced with pairs of GLY residues

All these double mutants that introduced additional hinges and removed charged residues lead to increased proportion of tilted conformations both in DLPC and DLPS membrane bilayers as compared to baseline counts of tilted and upright conformations.

#### I.4.3. Glycine-scanning mutants of TF LP

These mutations generate additional hinges in LP, similar to the double mutants described in the previous section. TF conformational changes in the TF_E213G mutant were similar for co-folding in the presence of DLPC and DLPS, in contrast to other mutations, when DLPC induced approximately two times more tilted conformations than DLPS and therefore, one can conclude that replacement of single Glycine had no considerable effect on conformation counts.

#### I.4.4 TF mutants that limit LP flexibility

TF_G211P mutant in the presence of DLPC and DLPS resulted in upright conformations, while there were two tilted conformations, suggesting that stiffening this hinge did not change the conformations in a phospholipid-dependent manner. By contrast, in the presence of DLPC, TF_G215P produced four tilted conformations out of ten in the output, whereas in the presence of DLPS, 100 % of the conformations were upright conformations. The mutant in which both G211 and G215 were replaced with valine residues produced 60 % upright and 40 % tilted (slow) conformations, while in the presence of DLPS, 90 % of conformations were active and 10 % tilted. Wild-type TF under these conditions produced 100 % active conformations. Thus, these mutations did not have considerable impact on phospholipid-dependent TF conformations.

**The TFfn2_C209S mutation**, which together with C186S mutation has previously been shown to inhibit TF activity by promoting an encrypted state of TF, resulted in 90 % upright structures corresponding to active TF conformation during co-folding in DLPC. In contrast, co-folding in the presence of DLPS resulted in 60 % of the upright and 40 % tilted TF conformations. These observations indicate that, in addition to the hinges in LP, the C209S mutation introduces flexibility that can affect the TF conformation. However, TF_C186S,C209S double mutant showed no conformational differences in the DLPC or DLPS co-folded structures. Thus, the role of Cys mutants in TF regulation, most likely, are mediated by the mechanism other than phospholipid induced TF conformations. Study of TF dimers described below (in section II of results and discussion) demonstrated that cholesterol induced monomerization of dimers was canceled by this mutant, that suggests that this mutant may have reduced activity, at least in part, due to stabilization of inactive TF dimers.

#### I.5.1. Direct contact between LP amino acid residues and phospholipid polar headgroups is important for TF conformation selection

Because both co-folding experiments and MD simulations confirmed the involvement of direct amino acid-phospholipid interactions in determining the TF conformation, we investigated these contacts in detail using the Pylipid software. This software uses interatomic distances to calculate phospholipid- amino acid interaction characteristics, including the interaction duration in the MD trajectory, lipid occupancy of the residues, and the number of contacting lipid molecules (21). Contacts are considered to exist between cutoff distances of 0.5 nm and 0.7 nm. The ten residues with the highest lipid occupancy for DLPC were 214 K, 216E, 217F, 218R, and 219E, while DLPS occupancy included the same residues as DLPC and 212Q. The remaining residues were from TFtmd and the proximal 181-185 loop of TF. MD validation of these results demonstrated 10 residues that showed the longest average interaction durations: 218R, 219E, 215G, and 216E. The remaining five residues were TFtmd residues that contacted DLPS because TFtmd was embedded in the membrane. Amino acid-lipid contacts in the case of “perpendicular to the membrane TFtmd included 218R and 219E, which were closest to TFtmd; other LP residues were not in contact with DLPS. Interestingly, TFfn2 remained tilted in this simulation. For the initially tilted TFfn2, which was straightened in the simulation, the main contacting residues were 218R and 217F included in the LP.

Analysis of the initial time steps of the ‘buried TFfn2’ MD simulation demonstrated contact between DLPS polar headgroups and LP residues. The longest contact and highest occupancy were found for R218 and K214, while residues with the largest number of surrounding lipids included K214, R218, TFtmd residues, and amino acid residues from the TFecd membrane proximal loops that potentially interact with the membrane bilayer.

#### I.5.2. Information theory measures Mutual Information (MI) and Transfer Entropy (TE) support the importance of LP dihedral angles in determining the TF conformation

To study the relationships between LP and the relative orientation of TF domain as well as the orientation of TFecd and TFtmd relative to the membrane bilayer plane, we used information-theoretic measures based on Shannon’s principles as implemented in the infomeasure python library. For strictly independent variables, MI is zero. For the dependent variables, MI is positive, increases with the strength of the relationship, and equals the entropy of a variable if it is a deterministic function of another. As seen from Fig. 4A, interdomain angles have strong dependence from LP dihedral angles MI values reaching 5 and higher.

**Figure 4.**
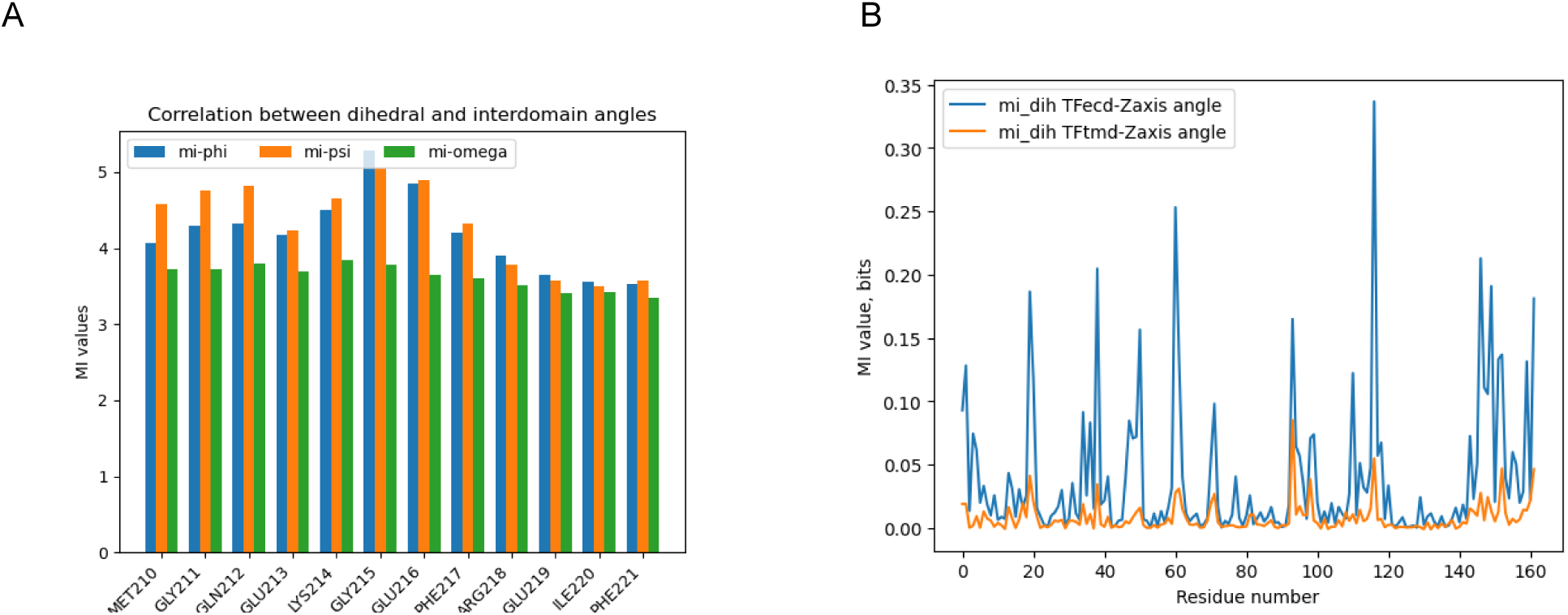
MI describing dependence between dihedral angles and interdomain angles. Panel A. MI values measured separately for phi, psi, and omega LP backbone dihedral angles in degrees and TFfn2-TFtmd interdomain angles for system containing PC (70 %) and PS (30 %). Panel B. MI between trigonometrically encrypted PHI-PSI angles (sin(phi)*sin(psi)) and angle between the principal axis of TFfn2 and Z-axis. Residue numbers 110-119 in this figure correspond to LP sequence G211- QEKGEFRE219.

Higher MI values are seen for residues of entire TFfn2-TFtmd fragment (Fig. 4B) for trigonometric encoding of phi and psi angles as well. However, this figure shows increased values not only for LP residues, but also for some additional residues. Because these additional residues were used in calculation of orientations of domains, these values should be disregarded during analysis.

TE measures the directed information flow between variables, in this case backbone dihedral and interdomain angles. TE is zero for independent variables or if the source does not improve the prediction of the destination (Fig. 5A and 5B). For the dependent variables, TE is a positive value that represents the reduction in uncertainty about the future of a destination variable provided by the past of the driver variable. Higher positive values of TE indicate a stronger causal influence or information flow from dihedral angles (source) to interdomain angles (target). TE from source to target does not necessarily equal TE from target to source. Fig. 5 demonstrates strong directed relationships in both direction for LP (residue numbers 110-120) pointing to possible causal link between these angles.

**Figure 5.**
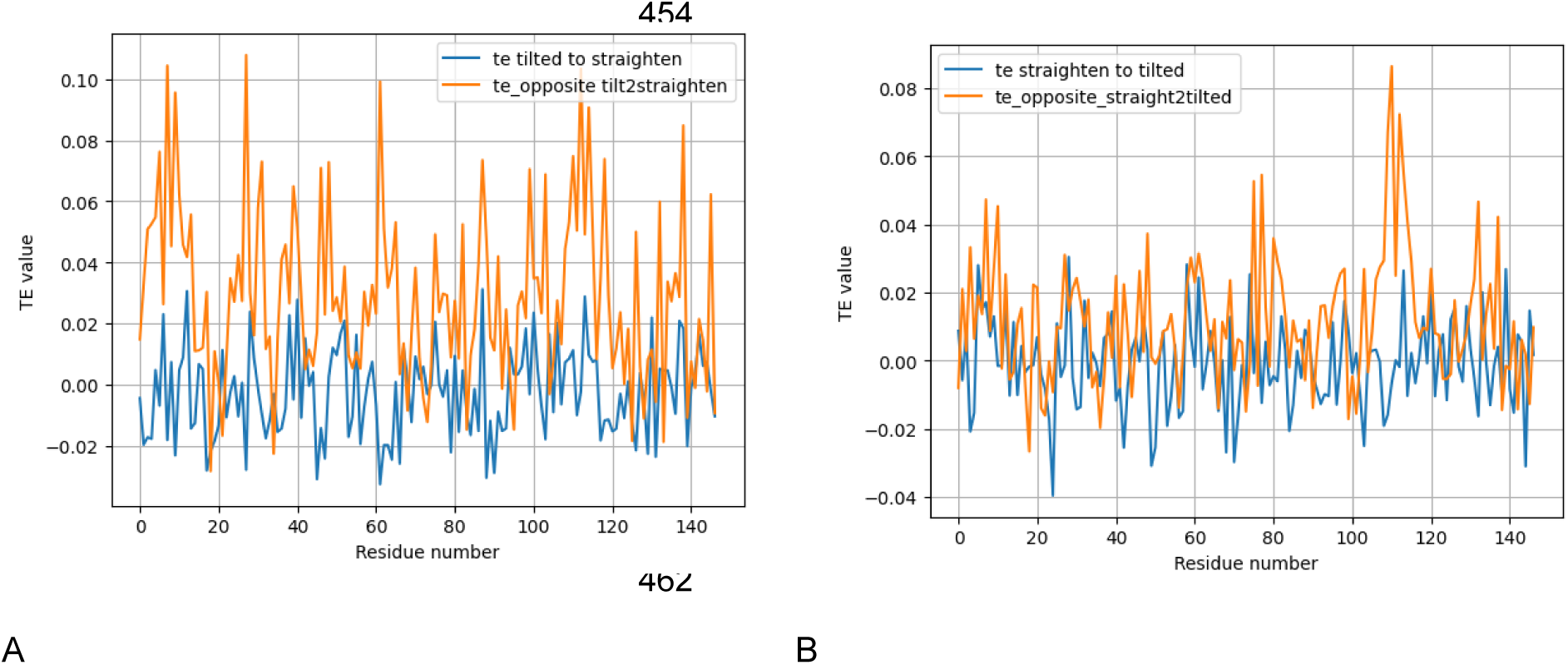
TE in direction from encrypted backbone dihedral angles to TFtmd-Z-axis angles and in opposite direction. Panel A. TE from dihedral to interdomain angles and vice versa for untilting of tilted TFfn2 (MD time steps 0-1000 or from 0 to 10 ns) Panel B. TE from dihedral to interdomain angles and vice versa for tilting of upright TF (MD time steps 1000-2300)

**Figure 6.**
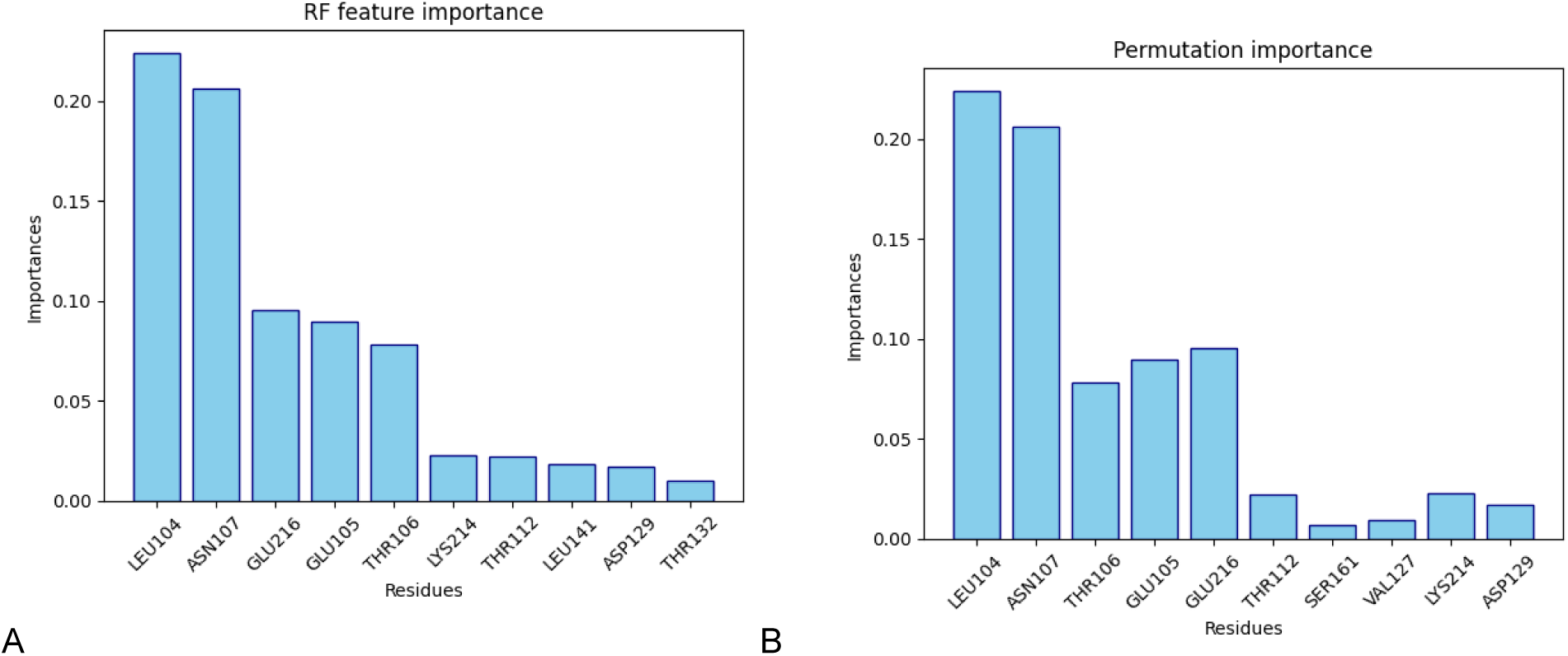
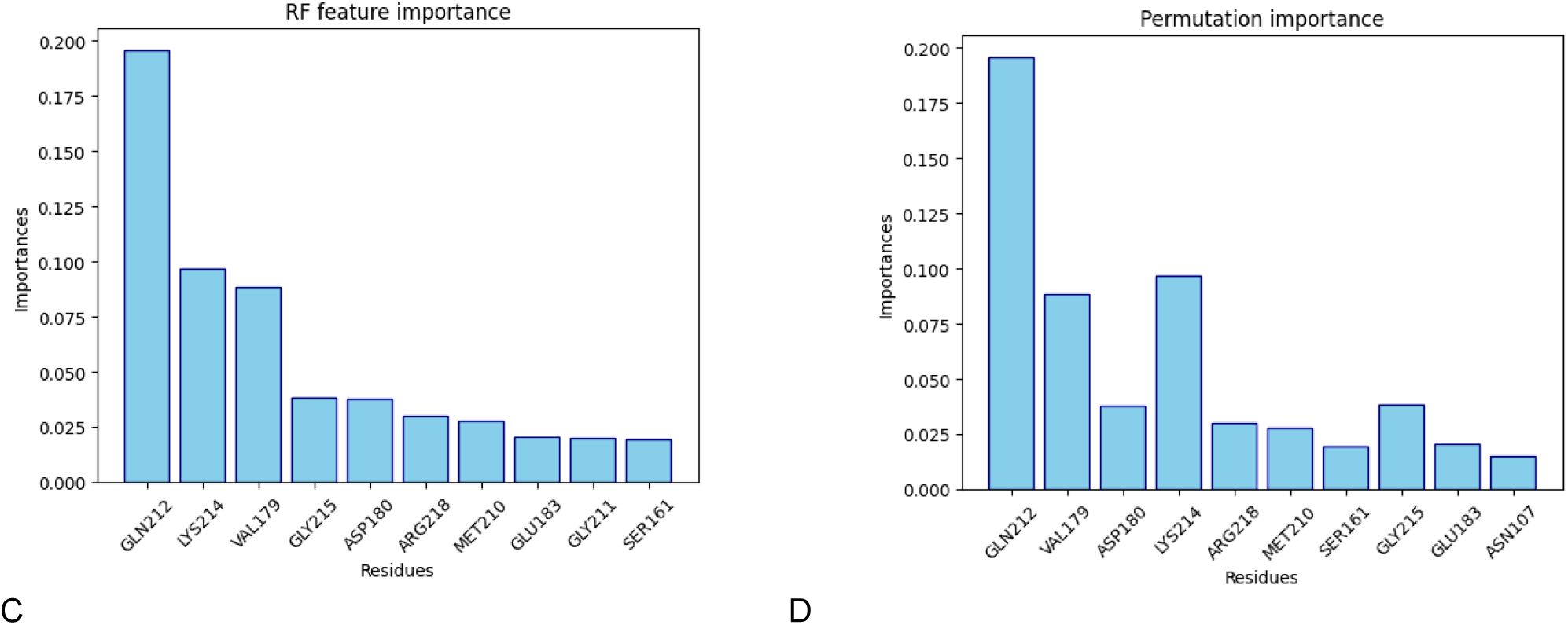
RF feature importance and permutation importance for the prediction of interdomain angles from the backbone dihedral angles. Panel A. RF regression feature importance and permutation importance (Panel B) for untilting of tilted TFfn2 in DLPS (MD steps 0-1000 from the trajectory of “buried” TFfn2-TFtmd simulation Panels C and D show the TF feature importance and permutation importance for MD frames 1000-2300, respectively.

#### I.5.3. RF regression and feature importance for LP residues

RF regression of the dataset composed of dihedral angles and interdomain angles demonstrated the importance of the Glu216 interaction with DLPS for untilting TFfn2. Tilting of upright TFfn2 could take place spontaneously, but MD-simulated dihedral angles show the importance of Q212, M210, K214, G211, G215, and R218.

#### I.5.4. Allosteric signaling in TFfn2-TFtmd

Analysis of cross-correlation matrices between residues in the TFtmd, LP, and distal FVIIa-binding domain of the TFecd allowed mapping of an allosteric pathway— a cascade of correlated movements—showing that lipid binding at the membrane interface transmits a structural signal through the LP to the distal from membrane FVIIa-binding site.

DCCM (Fig. 7) demonstrated a strong positive correlation between LP residues and membrane-proximal (phospholipid contacting) loops of TF and negative correlations for distal (relative to membrane) N- terminal loops of TF. The DCCMs compared in Fig. 7 represent an MD trajectory analysis for DLPS interacting (contacting) the LP amino acid residues (panel A) and a trajectory for MD simulation when LP residues had fewer contacts with DLPS molecules (panel B). Although the overall pattern of correlations is similar, there are considerable differences in the values of the correlation coefficients. The motions of the membrane-proximal loop atoms of TF were positively correlated with the motion of atoms in LP, whereas the atoms of the distal N-terminal loops were negatively correlated with the motions of LP atoms.

**Figure 7.**
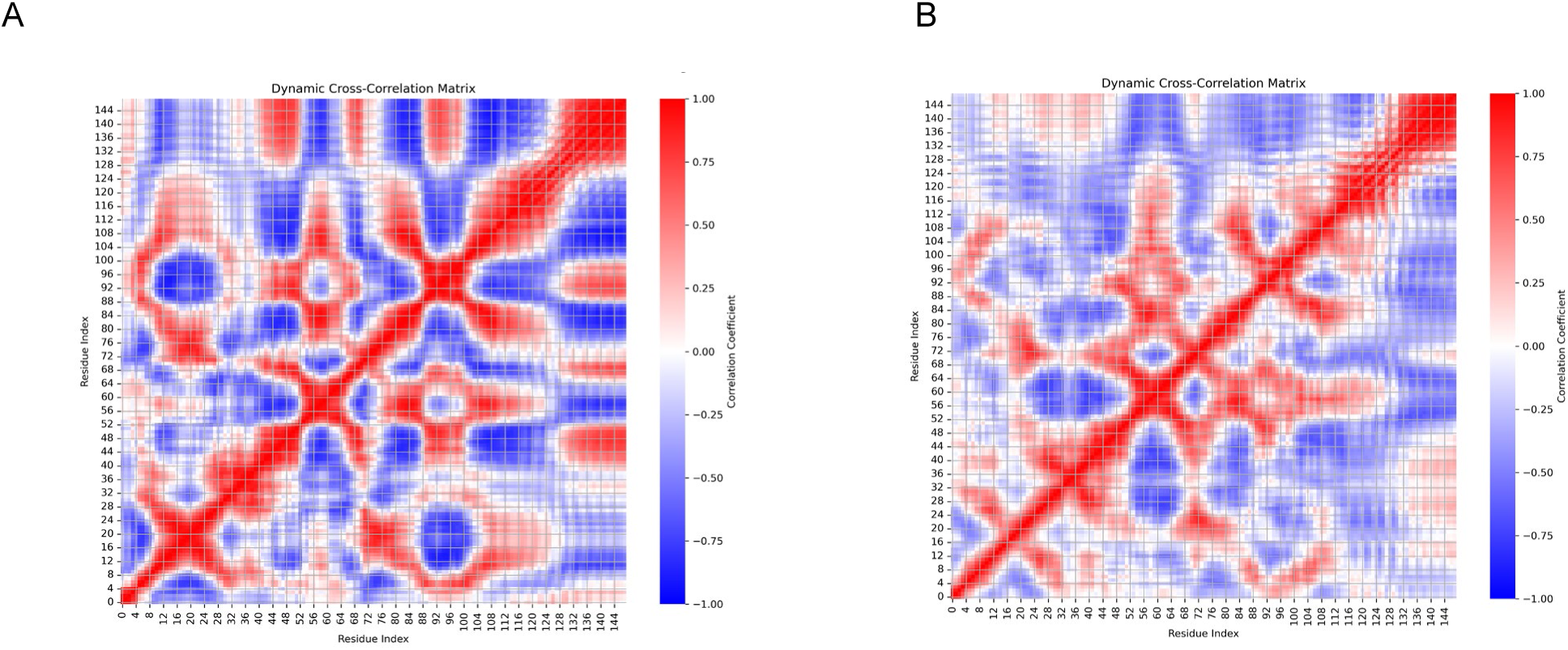
Dynamic cross-correlation matrix (DCCM). Value +1.0 (red) represents perfectly correlated motion. The atoms move simultaneously in the same direction. Value -1.0 (Blue) represents perfectly anti-correlated motion. When atom A moves up, atom B moves downwards. Value 0 (white) indicates Uncorrelated motion.

### I.6. Calculation of the Conformational Free Energy Landscape (FEL) for TFfn2-TFtmd during MD simulation

Conformational FEL were calculated using PCA by reducing high-dimensional MD simulation data into a 2D subspace, defined by the first two principal components (PC1 and PC2) calculated on atomic coordinates.

The FEL shown in Fig. 8 was constructed by plotting the free energy, computed from the probability distribution P (PC1, PC2), by converting probability distribution into free energy. This approach identifies stable low-energy conformers (blue and green regions) and their transition barriers. This is useful for analyzing conformational changes and protein flexibility. The results shown in Fig. 8 demonstrate difference in energy barriers for conformational changes due to LP-DLPS contacts. FEL calculated using LP dihedral angles also demonstrated different energy barriers for conformational transitions that depend on LP amino acid residues contacts with the polar headgroups of DLPS.

**Figure 8.**
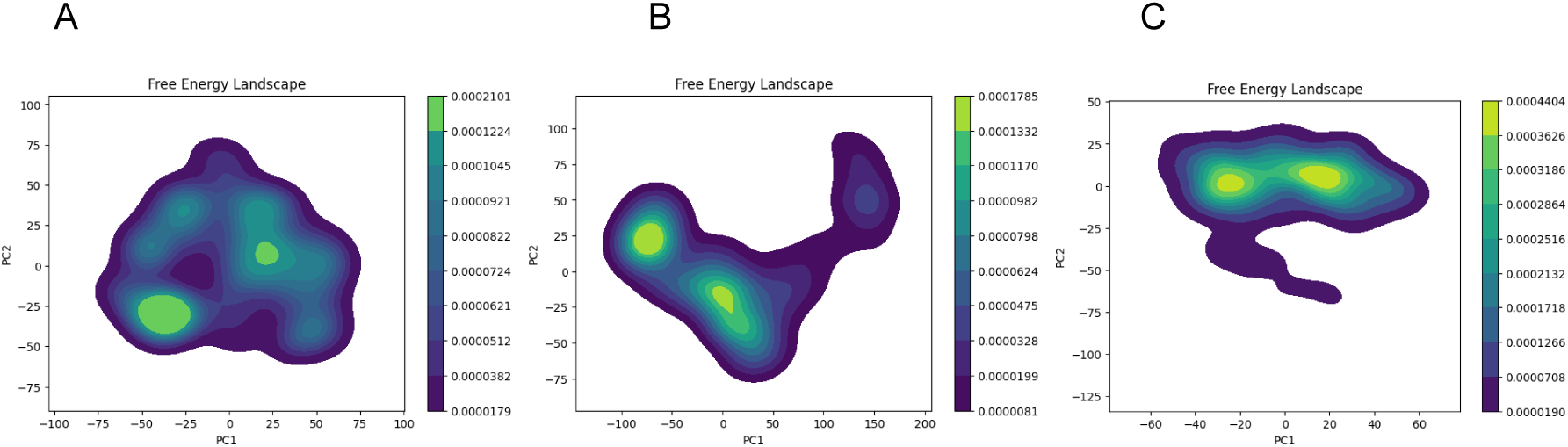
Free energy landscape for three trajectories of TFfn2_TFtmd in DLPS bilayer. PC1 and PC2 the first two principal components usually capture the largest and most significant conformational fluctuations. Energy Basins (deep blue-colored basins) represent stable, long-lived conformational states. Transition Paths (red/yellow) ridges and saddle points indicate energy barriers between states. Panel A. MD simulation of tilted TFfn2-TFtmd with LP-DLPS contacts. Panel B. Simulation of TFfn2-TFtmd without LP-DLPS contacts. Panel C. MD simulation of TFfn2-TFtmd with LP-DLPS contacts at the beginning of simulation.

#### II. SELF-REGULATION OF TF THROUGH OLIGOMERIZATION

It has been experimentally established that part of the TF antigen in the cell membrane can be detected in the form of dimers, and a smaller proportion of TF can be detected as higher-order oligomers (5, 6, 22). Because the experimental structure of the full-length TF dimer is unknown, we investigated the conformational ensemble formed by two TF sequences during co-folding alone or in the presence of different phospholipids using AF3 and Boltz2 software. Four types of dimers were detected in the conformational ensembles. We classified these dimers based on the relative position of Cys209 on the two TF monomers, considering Cys209 as a marker of the FVII binding surface of TF. The first type of dimer was denoted as a “FVII inside” type because Cys209 from each monomer was close to each other, and the bound FVII was located between the two TF molecules ( Fig. 10A). The second type was denoted as “FVII outside” because two bound FVII molecules enclose two TF molecules (Fig. 10C), and Cys209 in the monomers was located at the farthest distance from each other, facing externally. The third type of dimer was denoted as “FVII side-by-side” dimer because Cys209 on the monomers faced opposite directions and TF-FVII were side-by-side (Fig. 10E). The fourth type was rarely detected dimers with Cys209 on both monomers directed in the same direction, and two TF-FVII complexes “parallel” to each other that could be separated by an imaginary plane (not shown). The proportion of tilted or upright TF conformations among dimers was similar to the tilted and upright conformations of monomers in DLPC or DLPS, while the FVII-bound TF conformation of dimers was practically always upright.

#### II.1. Amino acid residues involved in the reversible dimerization of TF

Folding of two sequences representing full-length TF using both standalone AF3 and Boltz2 software or the AF3 web server resulted in the formation of dimers with contacting TFtmd and juxtamembrane cytoplasmic domain (Cys245) and extracellular domain LP-related residues (Fig. 9A). The TF dimers formed in the presence of DLPC, DLPS, and other phospholipids were similar to those formed in the absence of phospholipids. The exception was approximately 10% of dimers with two TFtmd forming a Y-like structure in which LP and neighboring residues in the TFtmd did not form contacts with the opposing monomer. The most frequent contacts for LP were GLN212, GLU213, LYS214, GLY215, GLU216, PHE217, and ARG218 with the same or neighboring residues of the opposite monomer. For TFtmd the most frequent contacts were between: ILE220, X, ILE224, X2, VAL227, X3, VAL231, X3, VAL235, X4, SER241, LEU242 in the monomers (Fig 9B, C, D). This sequence profile is different from known transmembrane domain interaction motifs, including the Val- based motif, despite the importance of the three Val residues for TFtmd dimerization. These contacts were found in the majority of TF dimers, and one could expect the formation of one main type of relative orientation of TFecd in dimers. However, the flexibility of LP enabled conformational freedom and different reorientations of TFecd for the same orientation of the TFtmd domains. Analysis of the conformational ensembles revealed at least four types of dimers distinguished by the relative orientation of TFecd (described in the previous section). Contacts between TFtmd residues located in the cytoplasmic leaflet of the membrane bilayer were consistently reproduced in dimers; as a result, the most frequent contact between monomers was mediated via Cys245 residues located in the intracellular domain of TF. Cys245 is palmitoylated in a fraction of TF in cells, and the de-palmitoylation of TF affects its function (23). A study of the co-folding of TF with CysP demonstrated a relative rotation of two TFtmd of TF monomers in dimers. In unmodified TF, Cys245 of the two monomers is positioned to allow disulfide bond formation. In addition to relative rotation, CysP allows the position of TFtmd to be adjusted to prevent hydrophobic disbalance. Consequently, palmitoylation affects the position of LP relative to the polar heads of phospholipids. Considering that LP functions as a lipid sensor that affects TF conformation, specifically the tilted or upright conformation of TFecd relative to the cell membrane surface. This is a plausible mechanism by which palmitoylation can affect TF function, related to TF translocation membrane microdomains with different lipid environments. Two palmitoylated Cys245 (CysP) from TF monomers were located within a distance that allowed indirect (via acyl chains) contacts between different monomers, thus contributing to the non-covalent binding of the two monomers. The Cys245 residue has previously been shown to form a disulfide bond that covalently links monomers (7, 17), while CysP is involved in the translocation of TF into cholesterol-sphingolipid-rich rafts in the cell membrane. Dimers with contacts at Cys245 have open TF FVII binding sites; therefore, these dimers are expected to be functionally active. These previously unknown TF conformations provide a structural basis for maintaining the encrypted and decrypted states of TF on the cell surface.

**Figure 9.**
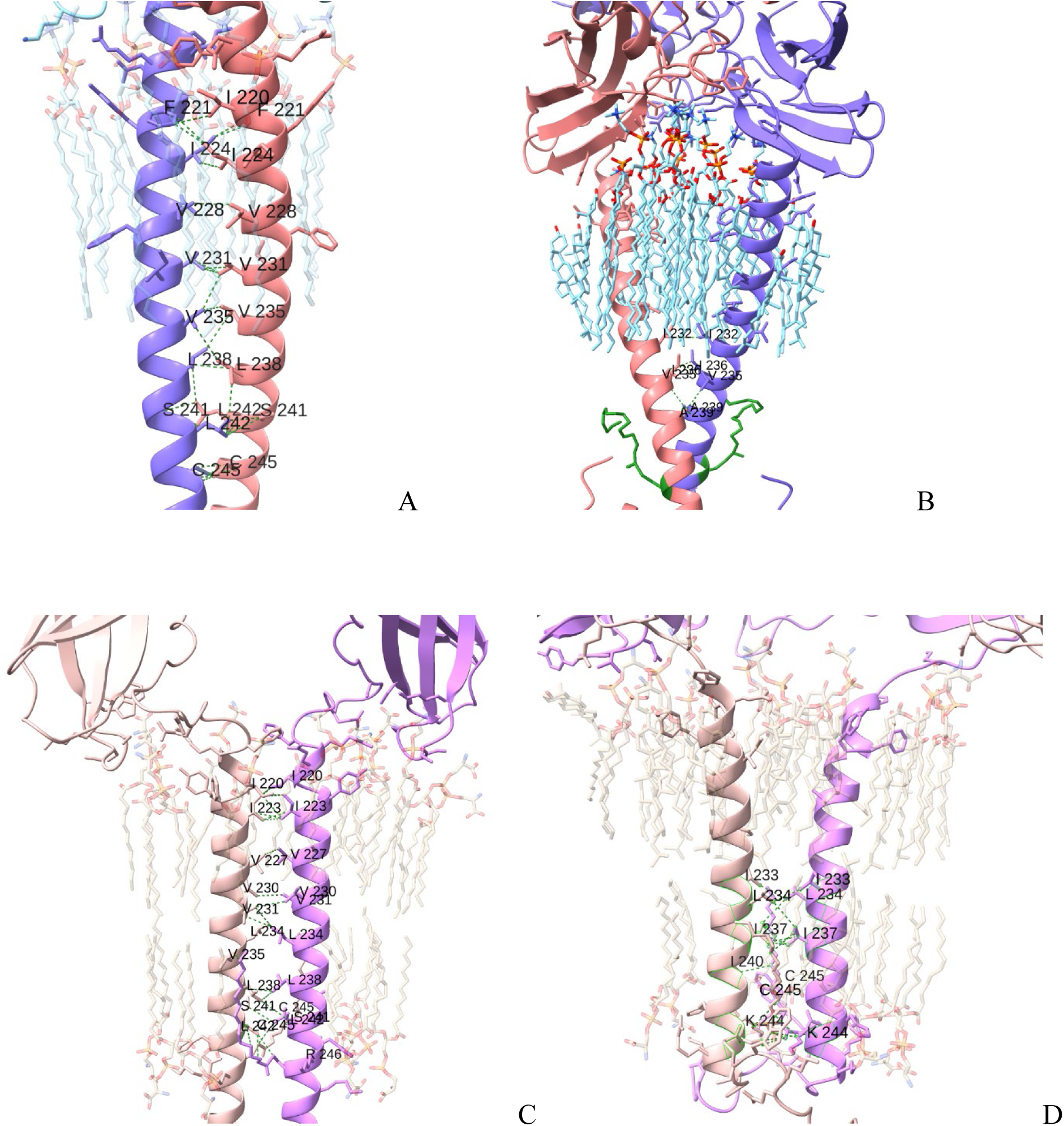
TFtmd contacts in dimers. TF monomers are shown in different colors, contacts are shown as dotted lines, contacting amino acid residues are labeled with 1-letter code and residue number in all panels. (A) TF dimer resulting from co-folding of two TF amino acid sequences in the presence of 20 phospholipid DLPC molecules. DLPC molecules are shown with 80% transparency. (B) TF dimer resulting from co-folding of two TF amino acid sequences in the presence of 20 phospholipid DLPC and 15 cholesterol molecules. Palmitoylated Cys245 is shown in green color. (C) TF dimer resulting from co-folding of two TF sequences in the presence of 20 phospholipid DLPS molecules. DLPS molecules are shown with 80% transparency. (D) TF dimer resulting from co-folding of two TF sequences in the presence of 20 phospholipid DLPS and 15 cholesterol molecules.

**Figure 10.**
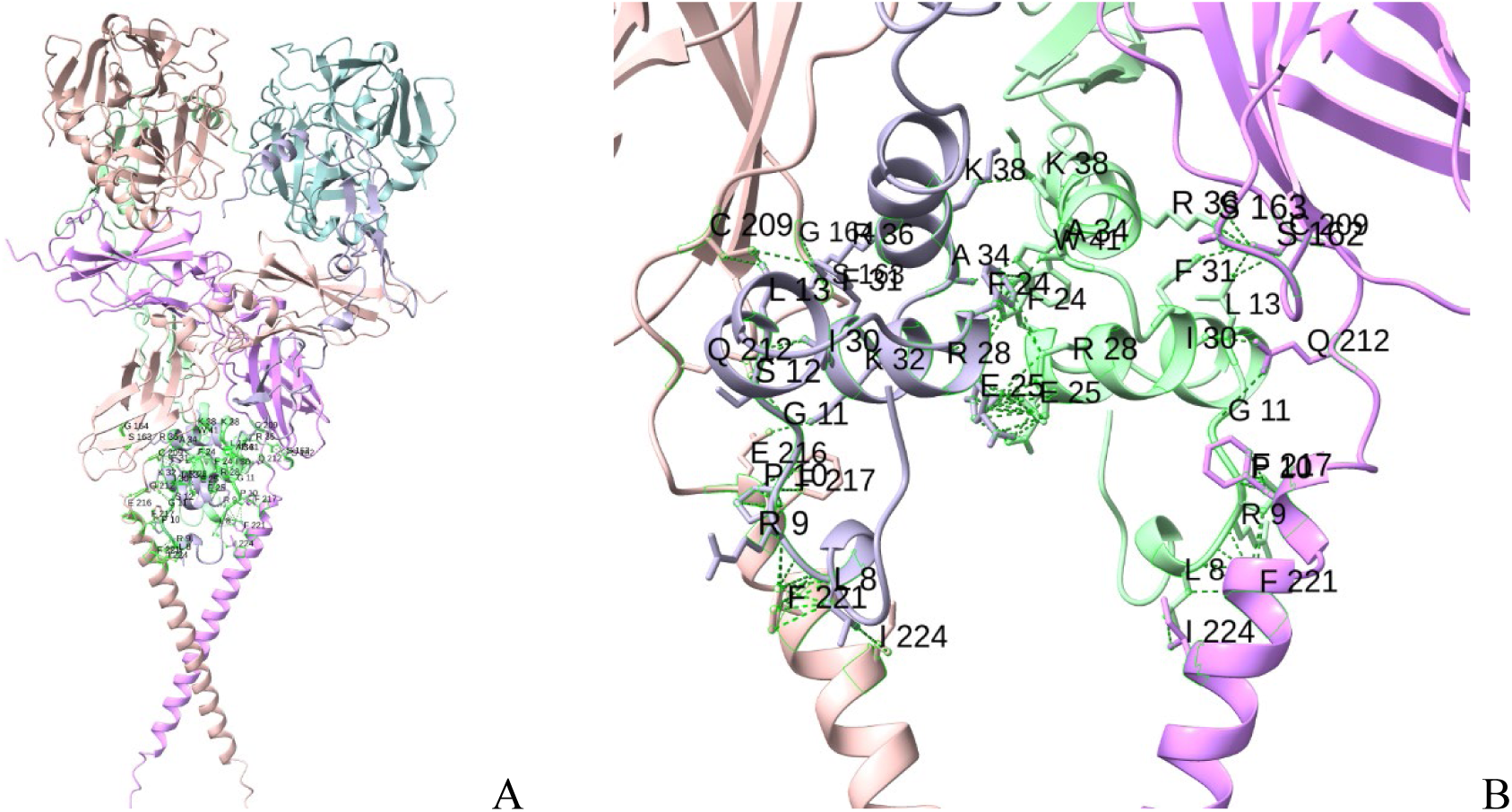
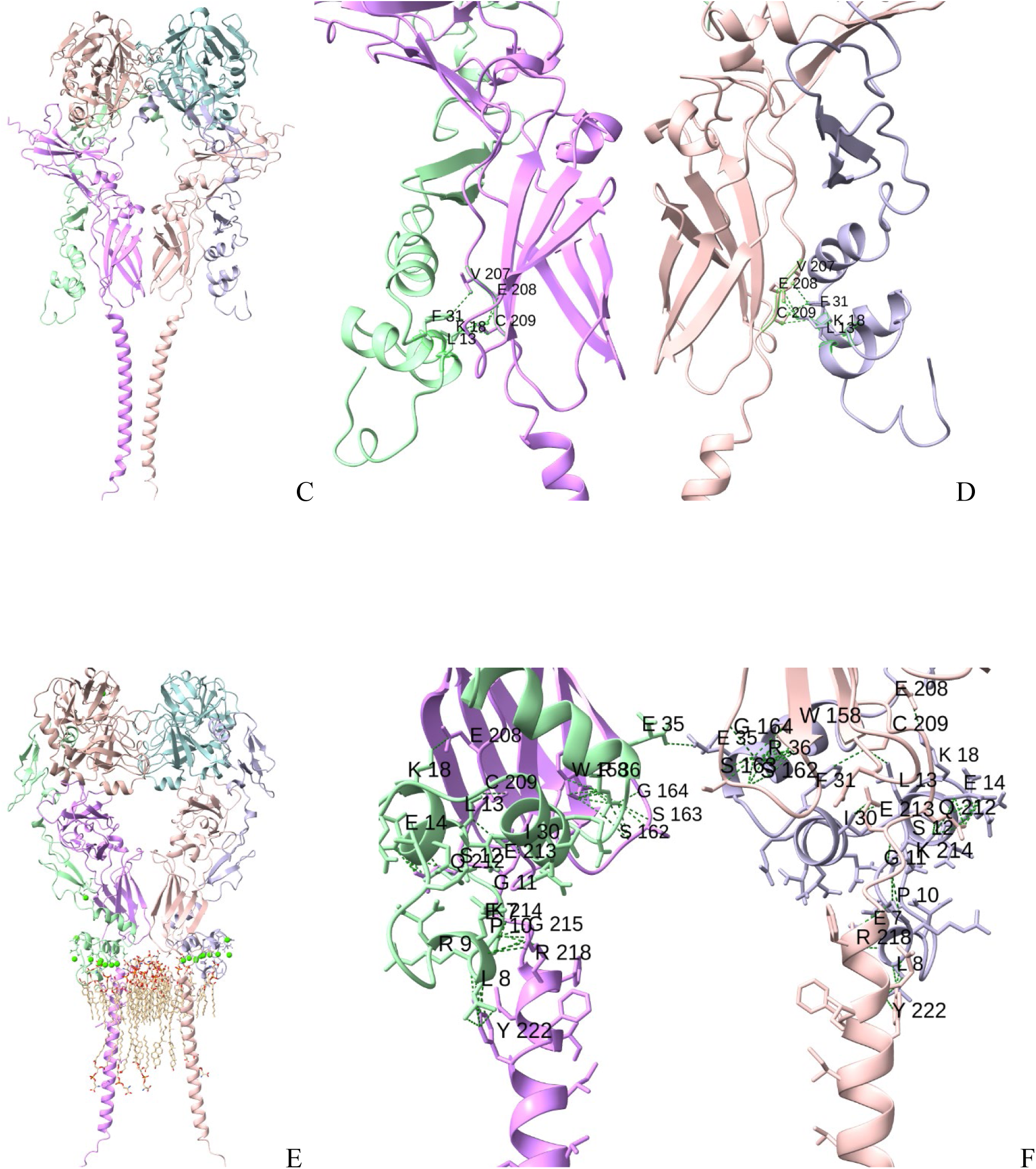
Three different types of complexes formed by TF dimer with two FVII molecules. TF monomers, FVIIa light and heavy chain are shown in different colors, contacts are shown as dotted lines, contacting amino acid residues are labeled with 1-letter code and residue number in all panels. (A) TF monomers enclose FVIIa molecules during co-folding of two TF and two FVIIa amino acid sequences. (B) Contacts between FVIIa Gla domain and TF LP amino acid residues in the complex shown in panel A (C) FVIIa molecules enclose TF monomers during co-folding of two TF and two FVIIa amino acid sequences. (D) Contacts between FVIIa Gla domain and TF LP amino acid residues in the complex shown in panel C (E) FVIIa molecules are located side-by-side relative to TF monomers during co-folding of two TF and two FVIIa amino acid sequences. (F) Contacts between FVIIa Gla domain and TF LP amino acid residues in the complex shown in panel E

**Figure 11.**
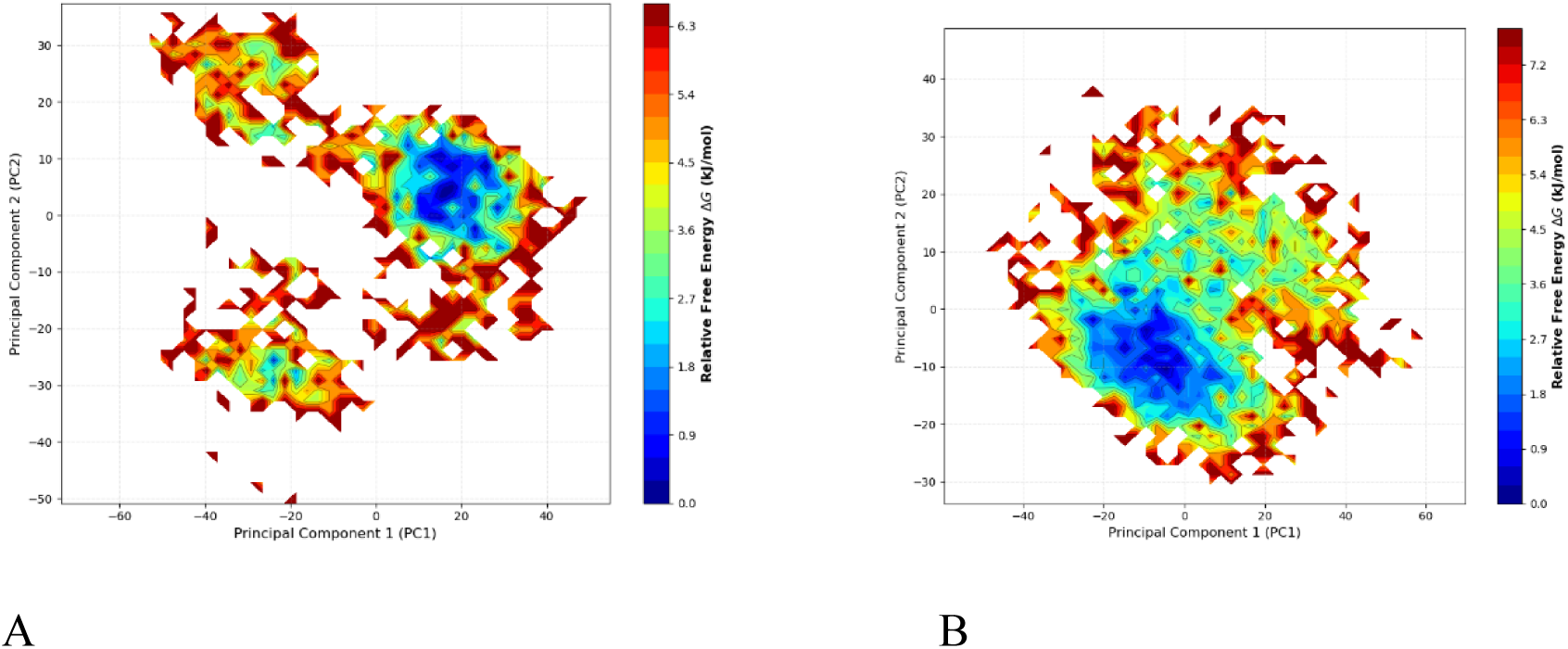
Free energy landscape of TF dimer. A. FEL in DLPS. B. FEL in DLPS:Cholesterol (4:3)

#### II.2. The conformational flexibility of the linker peptide enables FVII binding to TF dimers with upright TFecd, including dimers with sterically hindered FVII binding sites

Co-folding studies have shown that the FVII binding surface of TF is sterically occluded in “VFII inside” type of dimers. However, two FVII molecules formed complexes with two TF molecules in these and other situations as long as TFecd was in an upright position relative to the membrane surface (Fig. 10). Analysis of these structures showed direct contact between FVII and LP residues, suggesting that FVII induces conformational changes in LP that allow it to accommodate FVII binding to TF dimers. The flexibility of the LP allows TFecd and TFtmd to act as autonomous structural units with a wide range of relative orientation distances between TFecd, forming a variety of TF conformations. Investigation of these TF conformations, together with the conformations of TF monomers, revealed two previously unrecognized mechanisms of TF self-regulation related to the maintenance of its encrypted or active states. The first mechanism that employs the tilting of the TFecd was described in the previous sections. The second mechanism is based on TF self-association, forming TF dimers, as described in this section. The effect of these two mechanisms of TF self-regulation is best understood if the TF distribution on the cell surface is considered. The functional activity of TF located in lipid rafts differs from that of TF located in other membrane microdomains (12, 13). In addition to membrane microdomains, asymmetry in lipid distribution affects TF function. The main difference in the phospholipid composition of the outer and inner leaflets of the membrane bilayer that affects TF activity is that the outer layer of cell membranes mostly contains phosphatidylcholine and sphingomyelin, while the inner layer contains phosphatidylethanolamine, phosphatidylserine, and phosphatidylinositol. TF activation, concentrated in cholesterol-rich lipid rafts as well as in other microdomains, coincides with anionic phospholipid (PS) externalization (5, 24, 25). Lipid rafts are enriched in sphingolipids and cholesterol and packed with long, saturated fatty acid tails, whereas non-raft regions contain more unsaturated phosphatidylcholine and phosphatidylethanolamine, which create a more fluid environment. In addition to the formation of TF dimers, we tested the formation of trimers, tetramers, and pentamers. The TF sequence alone folded into oligomers via TFtmd contacts with additional contacts within the juxtamembrane regions of TF, including cytosolic Cys245 and some extracellular residues located in the LP-peptide. Co-folding of two, three, or four copies of the TF sequence in the presence of DLPS, DLPC, or phospholipids containing unsaturated fatty acid chains also resulted in the formation of TF oligomers held together by TFtmd contacts. These results suggest that the formation of a dimer prevents FVII binding to TF only for a proportion of dimers, most likely in dimers where TFecd is tilted and the TF FVII binding site is occluded. Thus, dimerization does not always prevent the binding of FVII to TF; it creates an additional tilted TF proportion of TF molecules that do not bind FVII or bind it slowly. The LP phospholipid polar head interactions in dimers were similar to those in TF monomers. Therefore, we observed more TF dimers with tilted conformations in the DLPC membranes than in the DLPS membranes. This picture was complicated in dimers because of the additional steric hindrance created by the proximity of TFecd in dimers. As a result, four types of TF dimers were formed in different proportions: DLPC co-folding resulted in approximately 50 % of “FVII inside” dimers with all tilted conformations, about 40 % “FVII side-by-side” and 10 % “FVII outside” conformations, all upright and capable of binding FVII. In DLPS co-folding experiments, approximately 30 % of the “FVII inside” and 10 % “FVII outside” conformations were all tilted conformations; largest proportion was about 60 % “FVII side-by-side” upright conformations. The LP plays an important role in forming the three types of dimers classified based on the relative orientation of the TFecd and described in previous sections. The TFtmd outer leaflet-located region and LP were responsible for the relative orientation of TFecd in dimers. The role of LP in the dimer type was even more evident in the complexes containing TF dimers and two FVII molecules. Amino acid residues directly interacted with FVII Gla-domain residues and changed their conformation to accommodate FVII bound to the TF dimer. Due to the flexibility of LP and its length (10 amino acid residues), all three described types of dimers were able to bind FVII after acquiring the LP conformation that allowed the creation of enough space between TF monomers to accommodate the FVII molecule bound to each TF monomer.

#### II.3. Cholesterol prevents the formation of TF dimers and maintains the monomeric state of TF

Co-folding of two TF sequences in the presence of DLPC, DLPS, and cholesterol prevented the formation of TF dimers. TFtmd was either completely separate from the closest TFtmd or, as shown in Fig. 9B, TF dimers held together by contacts at Cys245, had TFecd and LP separated by a large enough distance to accommodate FVII molecules. This observation was true for all TF oligomers, not just dimers. This provides a plausible explanation for the reduced FVII binding to cell surface TF after methyl-b- cyclodextrin treatment of cells (26). Cholesterol-dependent monomerization of RF suggests that the removal of cholesterol from lipid rafts most likely resulted in an increased number of dimers that were unable to bind FVII. This reduction in FVII binding was reversed after incubating the cells with methyl-b- cyclodextrin preloaded with cholesterol. This suggests that membrane cholesterol maintains the monomeric and active conformation of TF. Interactions of DLPC, DLPS, and cholesterol with TFtmd and juxtamembrane regions of TF. Co-folding of the TF sequence in the presence of different lipids demonstrated that DLPC mostly binds to the TFtmd region located in the outer leaflet of the membrane bilayer, while DLPS binds equally to both the outer and inner leaflet-related regions of TFtmd. The affinity of these phospholipids for TFtmd, as determined using Boltz2, was in the micromolar range. Cholesterol molecules bind to TFtmd with similar micromolar affinities, with some preference for the outer leaflet- related region of TFtmd. It was evident that the sterol part of cholesterol interacts with Phe217 of TF located in the LP; however, there was no cholesterol-TF interaction that could be described as a binding site for cholesterol. Interestingly, the cholesterol-induced monomerization of TF dimers was less prominent for TF mutant dimers in which Cys186 and Cys209 were replaced with Ser. This mutant is of interest because the disulfide bond between Cys186 and Cys209 has been debated as an allosteric bond that contributes to TF encryption/decryption, although the mechanism is still debated. TF Cys mutants were prone to forming dimers, and dimerization was not affected by cholesterol. There was some discrepancy between the AF3 and Boltz2 co-folding results in these experiments. AF3 co-folding experiments generated 100% dimerized TF mutants in the presence of cholesterol, whereas Boltz2 experiments showed a proportion of mutant TF as monomers in the presence of cholesterol. The higher proportion of inactive dimers formed from mutant TF may, at least in part, explain the reduced TF activity of Cys mutants of TF.

#### II.4. Comparison of Free Energy Landscape of TF dimer in DLPS with TF dimer in DLPS:Cholesterol

The FEL derived from the MD simulation of TF dimers in the presence of DLPS compared to the TF dimer in the membrane bilayer containing DLPS and cholesterol showed a clear difference (Figure 13) in FEL. In the presence of cholesterol, the FEL shows one energy minimum, whereas there are at least two energy minima. This corresponds to the stability of TF monomers surrounded by DLPS and cholesterol compared to the stability of the TF dimer in the presence of DLPS alone.

## Conclusion

A study of the conformational ensembles of TF and its dimer generated in the presence of different phospholipids revealed two mechanisms of TF self-regulation involved in maintaining the encrypted and decrypted functional states of TF. The first mechanism of TF self-regulation suggests that the tilted TF conformation may represent an encrypted (inactive) TF structure. In these structures, the principal axis of TFecd can reach from perpendicular to the membrane surface (active TF with upright conformation) to parallel (inactive TF). TFecd tilting creates steric hindrance for FVIIa binding to TF, especially when the FVII binding surface of TFecd faces the membrane surface. The conformational change between the tilted and upright conformations of TF was reversible, fast, and dependent on the phospholipid environment. The relative orientations of TFecd and TFtmd were determined by the interaction between the amino acid residues of LP and the polar heads of the membrane phospholipids. TF LP can be considered a phospholipid sensor that influences TF conformation by interacting with the polar heads of the surrounding phospholipids. The proportion of tilted TF conformations affects the rate of FVII binding to TF because the upright conformation is the optimal FVII binding conformation, whereas binding to tilted TF molecules with an obscured binding surface requires conversion of the tilted to upright conformation. This conformational equilibrium may explain the time-dependent feature of TF encryption, which demonstrated that TF that binds FVII within the first several minutes of incubation with the cells can express almost all cell surface TF activity (27). Moreover, the fact that a larger proportion of TF monomers acquire an upright conformation during co-folding in the presence of PS, as compared to a higher proportion of tilted conformations in the presence of phosphatidylcholine, can, at least in part, explain the decryption (activation) of TF by exposed PS (19). A tilted TF conformation is the prevailing conformation in all cellular microdomains before PS externalization because the main phospholipids in the outer layer of the membrane are PC and sphingolipids. This correlates with the encrypted state of TF in resting cells. The second mechanism of TF self-regulation employs non-covalent dimerization of TF, dependent on TFtmd and juxtamembrane amino acid residues of TF. At least four types of dimers can be distinguished using the relative orientation of the TFecd in the dimer structure. The activation sites of the two FVII molecules bound to TF in most dimers were positioned such that they allowed mutual activation of FVII. However, dimerization can reduce FVII binding to TF by 30-50 %, suggesting that a proportion of dimers are unable to bind FVII. This was evident when we compared the effect of cholesterol on TF dimers, which increased the proportion of active monomeric TF molecules, with our previous results that demonstrated reduced FVII binding to cellular TF after the removal of cholesterol.

## METHODS

### Materials

TF, its fragments, and mutants used in this study.

**Full length (flTF)** include amino acids 1-263 representing a mature form of the canonical TF: SGTTNTVAAYNLTWKSTNFKTILEWEPKPVNQVYTVQISTKSGDWKSKCFYTTDTECDLTDEIVKDVKQTYLARVFSYPAGNVESTGSAGEPLYENSPEFTPYLETNLGQPTIQSFEQVGTKVNVTVEDERTLVRRNNTFLSLRDVFGKDLIYTLYYWKSSSSGKKTAKTNTNEFLIDVDKGENYCFSVQAVIPSRTVNRKSTDSPVEC- 210-MGQEKGEFRE-220-IFYIIGAVVFVVIILVIILAISL-242-HKCRKAGVGQSWKENSPLNVS that includes a flexible linker (LP) (residues 210-219) connecting TFecd with TFtmd (residues 220-242). The membrane-proximal fragment of TF, amino acid residues102-263 include the second fibronectin type III- like domain TFfn2, LP, TFtmd, and TFicd. In some experiments parts of TFecd or TFicd sequence were removed as described in the results.

### TF LP mutants

1. The eight-alanine mutant: 212-AAAAAAAA-220 was used to test the role of the secondary structure and charged residues in phospholipid sensing.
2. Five double mutants of LP where charged amino acids and their neighbors were replaced with two glycine residues: p.C209G;p.M210G, p.Q212G;p.E213G, p.E213G;p.K214G, p.E216G;p.F217G, p.R218G;p.E219G.
3. Glycine scanning mutagenesis of LP to test the role of increased flexibility and charges: p.C209G, p.M210G, p.Q212G, p.E213G, p.K214G, p.E216G, p.F217G, p.R218G, and p.E219G
4. Hinge blocking mutants of LP where Gly was replaced with Pro or Val to test the role of limited hinge motions: p.G211P, p.G215P, and both p.G211V and p.G215V

**The phospholipids used in this study** (name, SMILES, or Chemical Component Dictionary (CCD) ID):

1. Three species of Phosphatidylcholine (PC): 1.1. DSPC (18:0/18:0) SMILES: CCCCCCCCCCCCCCCCCC(=O)OC[C@H](COP(=O)([O-])OCC[N+](C)(C)C)OC(=O)CCCC CCCCCCCCCCCCC. 1.2. DLPC (12:0 PC), 1,2-dilauroyl-sn-glycero-3-phosphocholine, SMILES: [O- ]P(OCCN+(C)C)(OCC@(OC(CCCCCCCCCCC)=O)COC(CCCCCCCCCC C)=O)=O. 1.3. CCD ID M2R PC with acyl chains (16/20)
2. Four species of Phosphatidylserine (PS) with acyl chain lengths of 6, 12, and 18 carbons: 2.1. DSPS (18:1-18:0 / 16:0-20:1), 2-amino-3-[[3-hexadecanoyloxy-2-[(Z)-icos-4-enoyl] oxypropoxy]- hydroxyphosphoryl]oxypropanoic acid;2-amino-3-[hydroxy-[2-octadecanoyloxy-3-[(Z)-octadec-4- enoyl]oxypropoxy]phosphoryl]oxypropanoic acid. SMILES: CCCCCCCCCCCCCCCCCC(=O)OC[C@H](COP(=O)(O)OC[C@@H](C(=O)O)N)OC(=O)CCCCCCCCC CCCCCCCC. 2.2. DLPS 1,2-Dilauroyl-sn-glycero-3-phospho-L-serine, saturated phospholipid (12:0 carbons). SMILES: CCCCCCCCCCCC(=O)OC[C@H](COP(=O) (O)OCC(N)C(=O)O)OC(=O)CCCCCCCCCCC. 2.3. CCD ID PSF (1,2-dicaproyl-sn-phosphatidyl-L-serine). SMILES: O=C(OC(COP(=O)(OCC(C(=O)O)N)O)COC(=O) CCCCC)CCCCC

### 2.4. CCD ID P5S, PS contains palmitoyl acid tails

3. Phosphatidylethanolamine (PE). DLPE 1,2-Dilauroyl-sn-glycero-3-PE. SMILES: O=C (OCC(OC(=O)CCCCCCCCCCC)COP(=O)(O)OCCN)CCCCCCCCCCC, CCD IDof PE: PTY

#### Other lipids

Phosphatidic acid (PA) as an example of a negatively charged phospholipid, Sphingomyelin, cholesterol (CCD ID CLR), glycolipid, palmitic acid, and linoleic acid.

#### Computational Modeling of TF Conformations

The flexibility of LP necessitates the use of methods capable of studying peptide-lipid interactions under conditions of changing peptide conformation. MD simulations are such an approach, which in this study revealed the phospholipid-dependence of TF conformations. Additionally, new artificial intelligence-driven approaches are emerging to model the full dynamic conformational ensembles of proteins and study protein folding in the presence of ligands and their influence. This study employed AlphaFold3 (AF3) (28–30) and Boltz2 (31) software packages to investigate effect of phospholipids on TF conformation mediated by the flexible peptide-phospholipid interactions. Using these two approaches, the interaction of flTF, fragments, or mutants of TF with phospholipids was investigated by allowing them to fold in the presence of ligands (termed co-folding).

MD simulations were used to validate the results of the co-folding studies, to elucidate the mechanism of lipid sensing by TF, and to reveal amino acid residues important for TF self-association. The input files for these studies were prepared using CHARMM-GUI Membrane Builder. The initial coordinates of the TF were generated by superposing the experimental structure of the TF (PDB ID 1dan) onto the predicted structures. TF was aligned such that its transmembrane domain was within the hydrophobic core of the bilayer using the OPM Database integrated with CHARMM-GUI (32). After visual verification that the TFtmd was aligned with the bilayer, in some experiments, the TF position was shifted relative to the membrane surface to study the role of lipid contacts with LP residues in lipid sensing. TF was simulated in DLPC or DLPS membrane bilayers with or without cholesterol that mimicked cell surfaces that effect activity of TF. Control experiments were conducted using phospholipids with longer acyl chains, as listed above, to delineate a possible role of the hydrophobic mismatch in selection of TF conformation. The system was solvated with TIP3P water model molecules. Counter ions Na^+^ or Cl^-^ were added to neutralize the total charge of the protein. Additional ions were added to achieve a physiological concentration of 0.15 M. The CHARMM36m force field (33), which is highly optimized for membrane systems, was used. Input topology and parameter files were generated for the NAMD MD engine using the CHARMM-GUI.

The simulation followed a multi-step protocol suggested in the CHARMM-GUI output: minimization followed by six steps of equilibration, where restraints on the protein and lipids were gradually released. The systems were equilibrated in the NVT ensemble in this protocol. The production run represented an all-atom simulation of the protein-membrane system in an NPT ensemble for 100 – 500 ns, as indicated in the results section.

Long-range electrostatic interactions in MD simulations were computed using the particle mesh Ewald summation. The SHAKE algorithm was used to restrain all carbon-hydrogen bonds (34, 35). The VdW nonbonded cutoff was set to 12 Å, with a smoothing cutoff of 10 Å. The saved trajectory files were analyzed using trajectory analysis tools provided by the VMD (36) software and complemented with other methods described in methods section.

Co-folding and MD experiments and analysis of the results were conducted on a Lambda workstation containing two NVIDIA GeForce RTX 3090 GPUs with 24 GB of GDDR6X VRAM memory each.

#### Datasets, analysis metrics, and software used in this study

Protein backbone dihedral angles: Phi and Psi dihedral angles of TF in the trajectory or separate molecules were determined using the MDAnalysis (37, 38) software. The angles between the principal axes of TFtmd and TFecd and between TFecd and the membrane plane (using the Z-axis as normal to the membrane) were calculated using a Python script created with help of Google Gemini (39) and dihedral angles were saved to disk as a NumPy arrays. These angles were used in the analysis directly as degrees or radians, or after trigonometric encoding of dihedral angles to generate one parameter equal to sin(phi)*sin(psi).

1. The tilt angle of TF was estimated as the angle between the TFecd principal axis relative to the membrane normal (Z-axis in the MD simulations) or the principal axis of TFtmd. Angles between 30-90 degrees were considered TF conformations that slowed down the association of FVIIa with TF. When the TF model and TF-FVIIa structure (PDB-ID 1dan) overlapped, creating clashes between membrane-proximal TF and FVIIa residues, the predicted structures were considered inactive. Thus, the TFecd tilt was used as a surrogate activity test to show conditions favorable for fast TF-FVIIa complex formation, conditions for binding at a slower rate, and inactive structures with clashes.
2. To analyze the link between the LP backbone dihedral angles and TFecd-TFtmd orientation angles, we used information-theoretical measures Mutual Information (MI) and Transfer Entropy (TE) calculated using the Infomeasure software (40). These data were complemented with Random Forest (RF) Regression analysis as implemented in scikit-learn (41), including both RF and Permutation feature importance analyses.
3. Allosteric Network Analysis: Dynamic cross-correlation matrices (DCCM) were used to visualize the correlation between the motions in the membrane proximal fragments of TF. Python scripts and functions to visualize DCCM as well as function to calculate and visualize Free Energy Landscapes (FEL) were created using function templates suggested by Google Gemini.
4. The FEL was determined using two collective variables: 4.1. Projections of MD trajectories onto Principal Components (PCA) to calculate the ΔG between different states of TF in trajectory, followed by the representation of FEL as a contour plot of PCA1 and PCA2 components. PCA was determined using the scikit-learn implementation. 4.2. FEL was also derived using phi and psi dihedral angles as collective variables and a Python script for the calculation of ΔG and plotting the data.

#### 5. Phospholipid – amino acid interactions were determined using the Pylipid (21) software, in addition to the visual analysis with VMD and ChimeraX (42) software

## RESOURCE AVAILABILITY

The resource availability section is required for all research articles. This section might also be required for applicable reviews and perspectives. This section must contain the following required subsections under the resource availability heading: "lead contact," "materials availability," and "data and code availability."

For complete information on requirements, including formatting instructions, refer to the Info for Authors page specific to the journal you are publishing with.

### Lead contact

• Requests for further information and resources should be directed to and will be fulfilled by the lead contact, Alexei Iakhiaev.

### Materials availability

• This study did not generate new unique reagents.

### Data and code availability

• A representative MD trajectory is available on Zenodo: Iakhiaev, Alexei (2026) Tissue Factor controls its own conformation via sensing the phospholipid environment [dataset]. Zenodo.https://doi.org/10.5281/zenodo.20345715

• Python scripts and other data are available upon reasonable request.

• Any additional information required to reanalyze the data reported in this paper is available from the lead contact upon request.

## ACKNOWLEDGMENTS

FUNDING SOURCES: NATIONAL SCIENCE FOUNDATION; AWARD NUMBER 2100878

## AUTHOR CONTRIBUTIONS

Conceptualization; methodology; investigation; writing – original draft; writing – review & editing; funding acquisition; resources; all was done by the only author of this manuscript, A.I.

## DECLARATION OF INTERESTS

A.I. declare no competing interests

## DECLARATION OF GENERATIVE AI AND AI-ASSISTED TECHNOLOGIES IN THE WRITING PROCESS

During the preparation of this work, the author used Paperpal in order to edit grammar. After using this tool or service, the author reviewed and edited the content as needed and takes full responsibility for the content of the publication.

## Notes

### Competing Interest Statement

The authors have declared no competing interest.

